# Effect of Aging, Sex, and Gene (Fbln5) on Arterial Stiffness of Mice: 20 Weeks Adult Fbln5-knockout Mice Have Older Arteries than 100 Weeks Wild-Type Mice

**DOI:** 10.1101/2023.05.30.542920

**Authors:** Hai Dong, Jacopo Ferruzzi, Minliang Liu, Luke P. Brewster, Bradley G. Leshnower, Rudolph L. Gleason

**Affiliations:** The Wallace H. Coulter Department of Biomedical Engineering Georgia Institute of Technology and Emory University, Atlanta, GA; The George W. Woodruff School of Mechanical Engineering Georgia Institute of Technology, Atlanta, GA; Department of Bioengineering The University of Texas at Dallas, Richardson, TX, United States; Emory University School of Medicine, Department of Surgery, Division of Vascular Surgery, Atlanta, Georgia; Atlanta VA Medical Center, Decatur, Georgia; Division of Cardiothoracic Surgery, Department of Surgery, Emory University School of Medicine, Atlanta, Georgia

**Author notes:** For correspondence: Rudolph L. Gleason Jr, Ph.D. The George W. Woodruff School of Mechanical Engineering Georgia Institute of Technology Technology Enterprise Park, Room 216 387 Technology Circle, Atlanta, GA 30313-2412. These authors contributed equally (Hai Dong, Jacopo Ferruzzi).

**Keywords:** Arterial stiffness, Fibulin-5 gene, Aging, Sex, Unified-fiber-distribution (UFD) model

## Abstract

The arterial stiffening is a strong independent predictor of cardiovascular risk and has been used to characterize the biological age of arteries (‘arterial age’). Here we revealed that the Fbln5 gene knockout (Fbln5^-/-^) significantly increases the arterial stiffening for both male and female mice. We also showed that the arterial stiffening increases with natural aging, but the stiffening effect of Fbln5^-/-^ is much more severe than aging. The arterial stiffening of 20 weeks old mice with Fbln5^-/-^ is much higher than that at 100 weeks in wild-type (Fbln5^+/+^) mice, which indicates that 20 weeks mice (equivalent to ∼26 years old humans) with Fbln5^-/-^ have older arteries than 100 weeks wild-type mice (equivalent to ∼77 years humans). Histological microstructure changes of elastic fibers in the arterial tissue elucidate the underlying mechanism of the increase of arterial stiffening due to Fbln5-knockout and aging. These findings provide new insights to reverse ‘arterial age’ due to abnormal mutations of Fbln5 gene and natural aging.

This work is based on a total of 128 biaxial testing samples of mouse arteries and our recently developed unified-fiber-distribution (UFD) model. The UFD model considers the fibers in the arterial tissue as a unified distribution, which is more physically consistent with the real fiber distribution of arterial tissues than the popular fiber-family-based models (e.g., the well-know Gasser-Ogden-Holzapfel [GOH] model) that separate the fiber distribution into several fiber families. Thus, the UFD model achieves better accuracies with less material parameters. To our best knowledge, the UFD model is the only existing accurate model that could capture the property/stiffness differences between different groups of the experimental data discussed here.

## Introduction

The cardiovascular risk can be predicted independently from arterial stiffness^1^. Vlachopoulos et al.^2^ showed that aortic stiffness is a strong predictor of future cardiovascular events and all-cause mortality. Arterial stiffening has been used to characterize the biological age of arteries (‘arterial age’)^3, 4^ and has also been recognized as a key initiator of many cardiovascular diseases^5^. For instance, arterial stiffening accelerates the pressure pulse wave velocity, which increases the afterload of the left ventricle, and ultimately promotes myocardial remodeling, dysfunction, and failure^6^. Thus, it is of great importance to understand how arterial stiffness/stiffening is affected by key factors, such as aging, sex, gene mutations, and physiological locations.

The protein Fibulin-5 (Fbln5) is an extracellular elastin-associated protein that connects to integrins and localizes tropoelastin to microfibrils as shown in the pioneering work by Yanagisawa et al.^7^ and Nakamura et al.^8^. Abnormal mutations of Fbln5 gene could results in cutis laxa syndrome in humans^9–11^. The Fbln5 deficiency alters arterial microstructures by disrupting the integrity of elastic lamellae^7, 8,12^, which results in elongated and tortuous central arteries. Aging is another major risk factor for cardiovascular diseases^3^. Prior work by Wan et al.^12,13^ and Ferruzzi et al.^14,15^ documented how aging and loss of elastic fiber integrity due to Fbln5 deficiency alters material properties across the central vasculature. However, it is still not fully understood how the Fbln5 and aging influence the intrinsic material stiffness/stiffening of arteries.

Unlike linear elastic materials (e.g., steel) of which the stiffness could be characterized by a single parameter such as Young’s modulus, there is no single simple parameter that can describe the stiffness of arterial tissues, due to their nonlinear and anisotropic mechanical behaviors^16–22^. Arterial tissues are comprised of networks of collagen/elastin fibers embedded in a ground matrix, and can be regarded as fiber-reinforced composites^23–28^. Our recently developed unified-fiber-distribution (UFD) model^29–31^ is a physical-structure-based model, which could accurately characterize the nonlinear and anisotropic mechanical responses of arterial tissues. The UFD model considers the fibers in the arterial tissue as a unified distribution, which is more physically consistent with the real fiber distribution of arterial tissues than the popular fiber-family-based models that separate the fiber distribution into several fiber families^23, 24, 32, 33^. Here, based on a total of 128 biaxial testing samples of mouse arteries^14, 15^ and the UFD model^29–31^, for the first time, we elucidated how the aging, sex, and Fbln5 gene affect the intrinsic arterial stiffness of mice at five different locations, including ascending thoracic aorta (ATA), descending thoracic aorta (DTA), suprarenal abdominal aorta (SAA), infrarenal abdominal aorta (IAA), and common carotid artery (CCA). To our best knowledge, the UFD model is the only existing accurate model that could capture the property/stiffness differences between different groups of the experimental data discussed here.

## Results

There are four parameters in the UFD model (for details see Eq. 1 in Methods Section), including the arterial matrix modulus (*c*), arterial fiber initial modulus (*k*_1_), arterial stiffening factor (*k*_2_), and fiber distribution-related parameter (ζ). These microstructural parameters of arterial matrix and fibers are difficult to measure directly from the experiments. We obtained the four parameters *c*, *k*_1_, *k*_2_, and ζ for each of the 128 arterial samples (see Methods Section), by fitting the cylindrical biaxial testing data (pressure, diameter, axial force, and axial stretch) during fixed-length inflation (three protocols, Fig. 1a) and fixed-pressure axial extension (four protocols, Fig. 1b) trajectories with the UFD model. Excellent fitting accuracy was obtained (Fig. 1a, b), with the relative error (defined in Eq. 3) to be *ER* = 5.09% ± 0.32% (mean±standard error) and the coefficient of determination to be *R*^2^ = 0.9363 ± 0.0032. The values of *c*, *k*_1_, *k*_2_, and ζ, as well as, *ER* and *R*^2^ for each of the 128 arterial samples based on the seven protocols were given in Extended Data Tables 1-5. To further validate the accuracy of the UFD model, we also fitted the parameters based on five of the seven protocols, and used the fitted parameters to predict the results of the other two protocols. The predicted results also agree very well with the experiments (see Extended Data Fig. 1).

**Fig 1.**
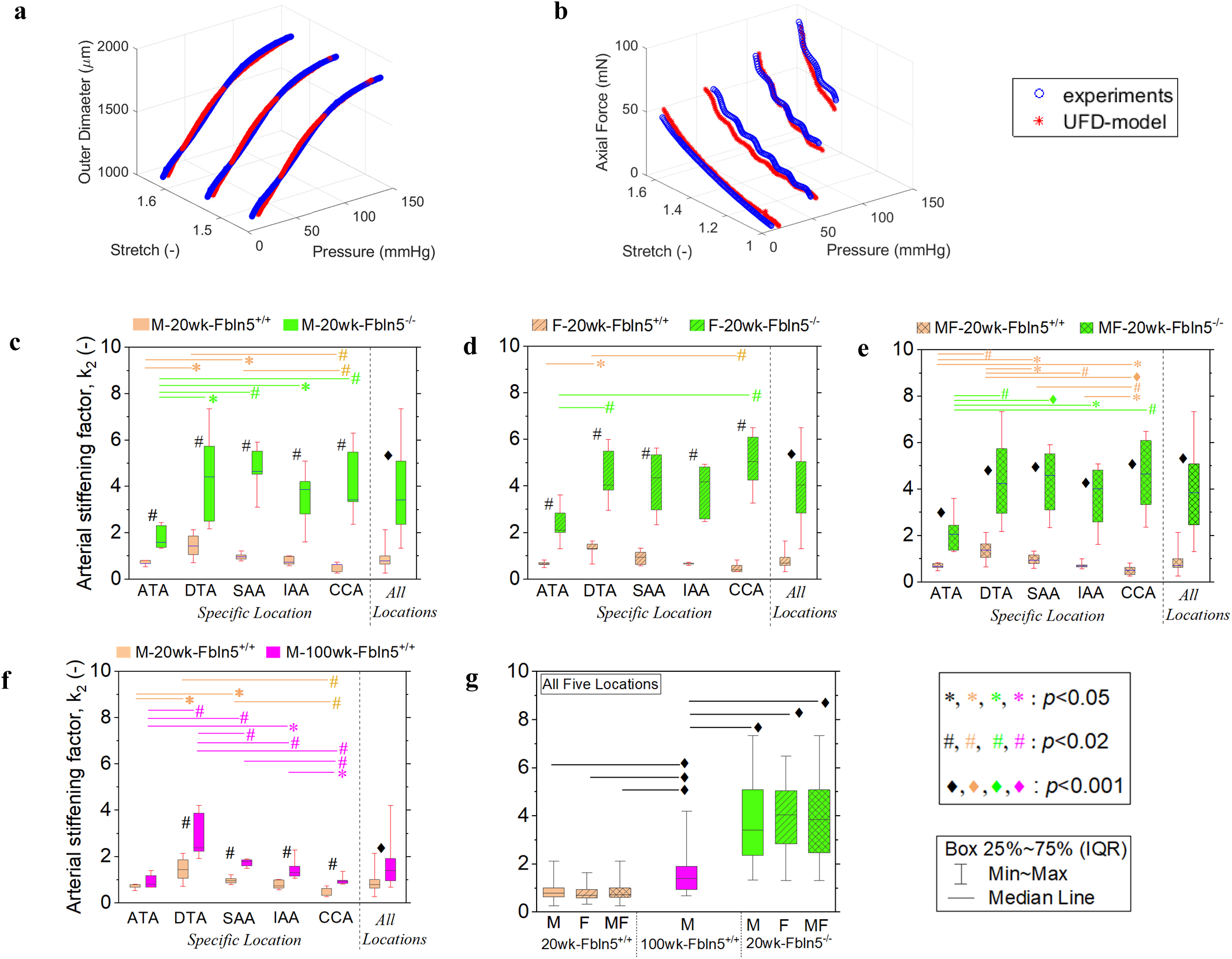
**a-b**: A representative example of the fitting results of Pressure-Diameter relation (**a**) and Pressure-Axial Force relation (**b**), based on the UFD model^6–8^. Excellent fitting accuracy was obtained, with the mean error for all samples (n=128) to be *ER* = 5.09% ± 0.32%, and *R*^2^ = 0.9363 ± 0.0032. **c-g:** Arterial stiffening factor (*k*_2_) of male (M) and female (F) mice at 20 weeks (20wk) and 100 weeks (100wk) old, for wild-type (Fbln5^+/+^) and knock-out type (Fbln5^-/-^). **g:** All five locations analyzed together. ATA: ascending thoracic aorta, DTA: descending thoracic aorta, SAA: suprarenal abdominal aorta, IAA: infrarenal abdominal aorta, and CCA: common carotid artery. MF: male and female analyzed together. *p-*vlaues were based on nonparametric Wilcoxon rank sum test. Asterisks 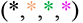: *p*<0.05, Hash signs 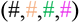: *p*<0.02, and Diamonds 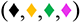: *p*<0.001. Black symbols (*, #, ♦) indicate statistic difference between different genotypes and/or different ages. Colored symbols 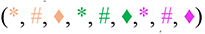 near the right end of a line indicate statistic difference between different locations. Arterial stiffening with Fbln5^-/-^ is significantly larger than that with Fbln5^+/+^, for both M and F, at each of the five locations (**c**, **d, e**). The aging effect also significantly increases the arterial stiffening (**f**). Arterial stiffening at 20 weeks old with Fbln5^-/-^ is even larger than that at 100 weeks old with Fbln5^+/+^ (**g**), indicating that, the 20 weeks old mice (equivalent to ∼26 years old human^35^) of Fbln5^-/-^ have a higher ‘arterial age’ than 100 weeks old wild-type mice (equivalent to ∼77 years old human^35^) of Fbln5^+/+^. Arterial stiffening factor follows the same pattern in terms of locations for M (**c**) and F (**d**), with either Fbln5^+/+^ or Fbln5^-/-^. For Fbln5^+/+^ groups (orange boxes in **c**, **d, e**), arterial stiffening factor increases from ATA to DTA, and then decreases to IAA to CCA; For Fbln5^-/-^ groups (green boxes in **c**, **d, e**), the arterial stiffening factor increases from ATA to DTA, and then keeps within the same range among DTA, SAA, IAA, and CCA.

We performed both parametric and nonparametric tests (two-sample t-test and Wilcoxon rank sum test, respectively, see Methods Section) to analyze the difference of the parameters from different groups. The two different tests yield similar results (presented either in Figs. 1-5 or in Extended Data Figs. 2-5).

### Arterial stiffening factor (*k*_2_) and histological microstructure of elastic fibers

The arterial stiffening (*k*_2_) of mice with Fbln5 gene knockout (Fbln5^-/-^) was significantly larger than that of the wild-type (Fbln5^+/+^), for both male (M) and female (F) when considering data of all five locations together (Fig. 1c for M: *p*<0.001, “*All Locations”* [n=25 for each Fbln5^-/-^ & Fbln5^+/+^ groups]; Fig. 1d for F: *p*<0.001, “*All Locations”* [n=25 for each genotype group]), as well as when considering data separately for each specific location (Fig. 1c, d: all *p*<0.02, each “*Specific Location”* [n=5 for each genotype group]). When pooling the male and female samples together, the *p-*values of the arterial stiffening between the Fbln5^-/-^ and Fbln5^+/+^ groups decreased at all five locations (Fig. 1e: all *p*<0.001 for each *Specific Location* [n=10 for each genotype group], and *All Locations* [n=50 for each group]).

The nonlinear mechanical responses (stiffening at large deformation such as greater than 5% strain) of arterial tissues are attributed to the progressive engagement of fibers under increases in strain. Our new histological results (first and third columns in Fig. 2) showed that the Fbln5-knockout caused fragmentation and waviness loss (increased wave length and decreased wave amplitude) of the elastic fibers at all five investigated locations, which is consistent with previously work^7, 8, 14^ and well explains the increase of arterial stiffening by the Fbln5-knockout, since the waviness loss results in earlier (in strain) engagement of the elastic fibers and makes the tissue stiffer at the same deformation. The micro-structure of collagen fibers is not affected much by the Fbln5-knockout^7, 8, 14^.

**Fig 2.**
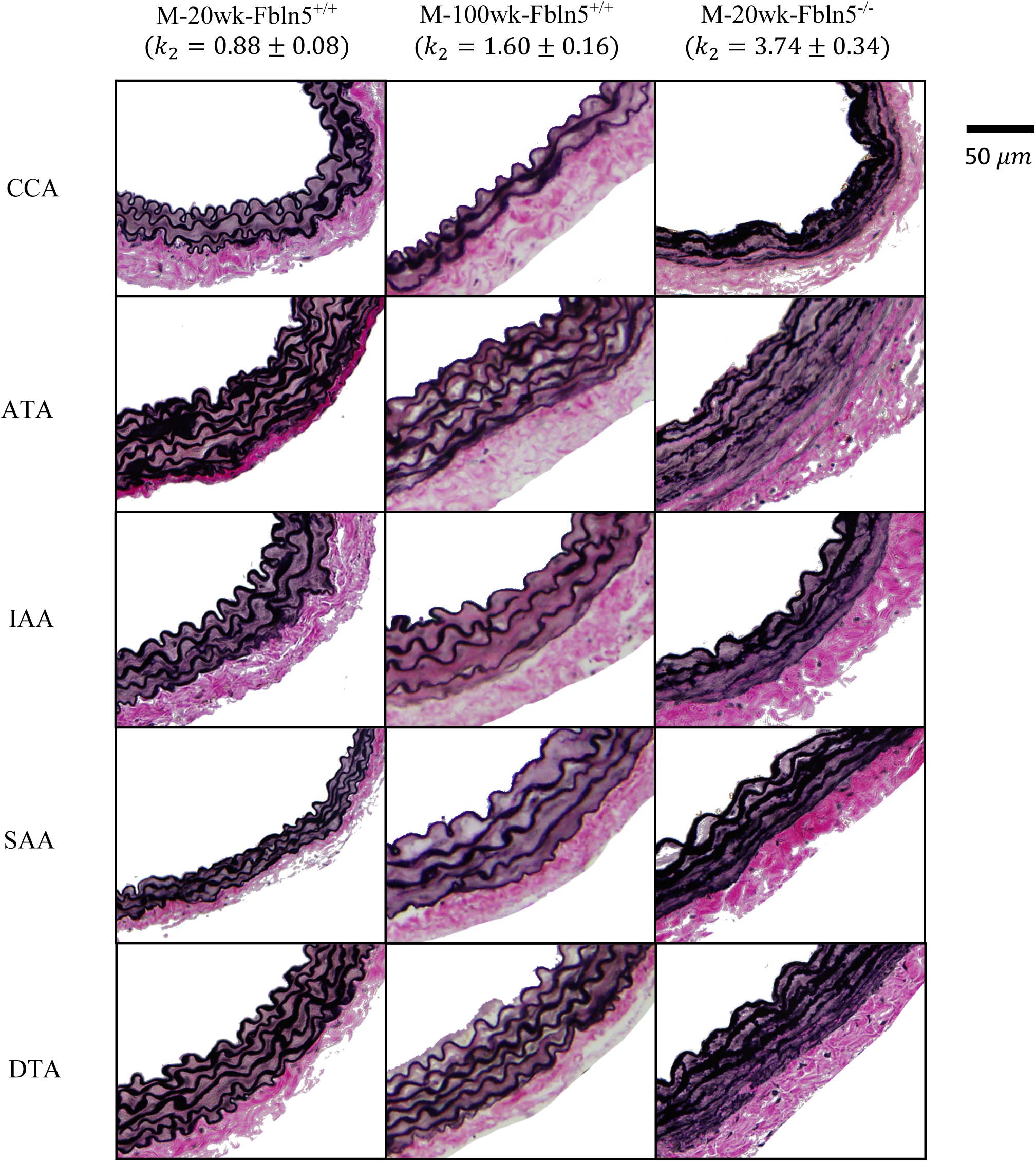
Elastic fibers of male mice at 20 weeks old (20wk, equivalent to ∼26 years old humans^35^) and 100 weeks old (100wk, equivalent to ∼77 years humans^35^), for wild-type (Fbln5^+/+^) and knockout type (Fbln5^-/-^). CCA: common carotid artery, ATA: ascending thoracic aorta, IAA: infrarenal abdominal aorta, SAA: suprarenal abdominal aorta, and DTA: descending thoracic aorta. The elastic fibers lose waviness (increased wave length and decreased wave amplitude) with both aging and Fbln5-knockout, at all five investigated locations, which well explains the increase of arterial stiffening by the Fbln5-knockout and aging. The Fbln5-knockout results in more severe waviness loss of the elastin fibers which is perfectly consistent with the results, independently obtained from the mechanical testing, that the arterial stiffening factor of 20wk-Fbln5^-/-^ mice is larger than that of 100wk-Fbln5^+/+^ mice.

Aging (20 vs 100 weeks old) significantly increased the arterial stiffening in wild-type mouse arteries (Fig. 1f: *p*<0.001, “*All Locations”* [n=25 for each 20 & 100 weeks groups]). As to the *Specific Location,* the arterial stiffening factor of mice at 100 weeks old was notably larger than that of mice at 20 weeks old at locations of DTA, SAA, IAA, and CCA (Fig. 1f: *p*<0.02, [n=5 for 20 & 100 weeks groups]). These findings could also be explained by the waviness loss of elastic fibers due to aging (first and second columns in Fig. 2). Moreover, the accumulation of the collagen fibers with aging^15, 34^ may also contribute to the increase of the arterial stiffening in aging 100 weeks old mice.

The mouse arterial stiffening of 20 weeks old with Fbln5^-/-^ was even larger than that of 100 weeks old with Fbln5^+/+^, as shown in Fig. 1g and Extended Data Fig 2e (all *p*<0.001, [n=25 for M or F groups, and n=50 for MF groups]). The biological age of arteries (‘arterial age’), usually characterized by the arterial stiffening^3, 4^, could deviate from the chronological age. Here the arterial stiffening factor (*k*_2_) can be regarded as a measurement of ‘arterial age’, which suggests that the 20 weeks old mice (equivalent to ∼26 years old human^35^) of Fbln5^-/-^ have a higher ‘arterial age’ than 100 weeks old wild-type mice (equivalent to ∼77 years old human^35^) of Fbln5^+/+^. The histological microstructures of elastic fibers (Fig. 2) show that the Fbln5-knockout results in more severe waviness loss than aging from 20 weeks to 100 weeks for all the five investigated locations, which explains the higher ‘arterial age’ of 20 weeks mice with Fbln5^-/-^ than 100 weeks mice with Fbln5^+/+^. These results are also consistent with the effect of Fbln5 mutations on the abnormal looseness and wrinkling of skins in both mice^7, 8^ and humans^11, 36, 37^ with Fbln5-related Cutis Laxa Syndrome. Thus, the knock-out of Fbln5 gene may be considered as a super-aging effect.

No statistical difference of the arterial stiffening was found between the male and female groups with the same genotype at any of the five locations. But it is interesting to note that the arterial stiffening across different locations along the vascular followed the same pattern for male (Fig. 1c) and female (Fig. 1d) mice with the same genotype (either Fbln5^+/+^ or Fbln5^-/-^). For the Fbln5^+/+^ groups (orange boxes in Fig. 1c, d), the arterial stiffening factor increased from ATA to DTA, and then decreased to IAA to CCA; For the Fbln5^-/-^ groups (green boxes in Fig. 1c, d), the arterial stiffening factor increased from ATA to DTA, and then kept about the same level among DTA, SAA, IAA, and CCA. When considering the male and female samples together, all the *p-* values between groups at different locations decreased for both Fbln5^+/+^ and Fbln5^-/-^ (Fig. 1e), which indicates that the conclusion of the same location pattern with the same genotype is strengthened with increasing sample size. The arterial stiffening may be determined by both the wave length and wave amplitude of the elastic fibers in Fig. 2, and the arterial stiffening *k*_2_ can be regarded as a measure of the average of the combined wave length and amplitude.

Comparison between Fig. 1c and Fig. 1f showed that the Fbln5 knockout changes the location patterns but the aging effect does not, despite that both aging and the Fbln5 knock-out increase the arterial stiffening factor, which suggests that the genotype is the key factor controlling the location patterns of arterial stiffening.

### Arterial matrix stiffness (*c*), circumferential fiber component (ζ), and fiber initial stiffness (*k*_1_)

The parameters *c*, *k*_1_ and ζ also followed similar patterns in terms of locations for both male and female mice with the same genotype, as shown in Figs. 3-5a, b, c, where the mean and standard error with the results of the parametric two-sample t-test were presented. The arterial matrix stiffness (*c*) decreased from ATA to SSA to CCA for Fbln5^+/+^ groups (orange bars in Fig. 3a, b, c), while first increased from ATA to SSA, and then decreased to CCA for Fbln5^-/-^ groups (green bars in Fig. 3a, b, c). The Fbln5 knockout showed decreases of the matrix stiffness at ATA (Fig. 3a, b, c), while then increases of the matrix stiffness at IAA and CCA (Fig. 3c). The aging effect has little effect on the matrix stiffness at most locations (only reduces the value at SAA in Fig. 3d).

**Fig 3.**
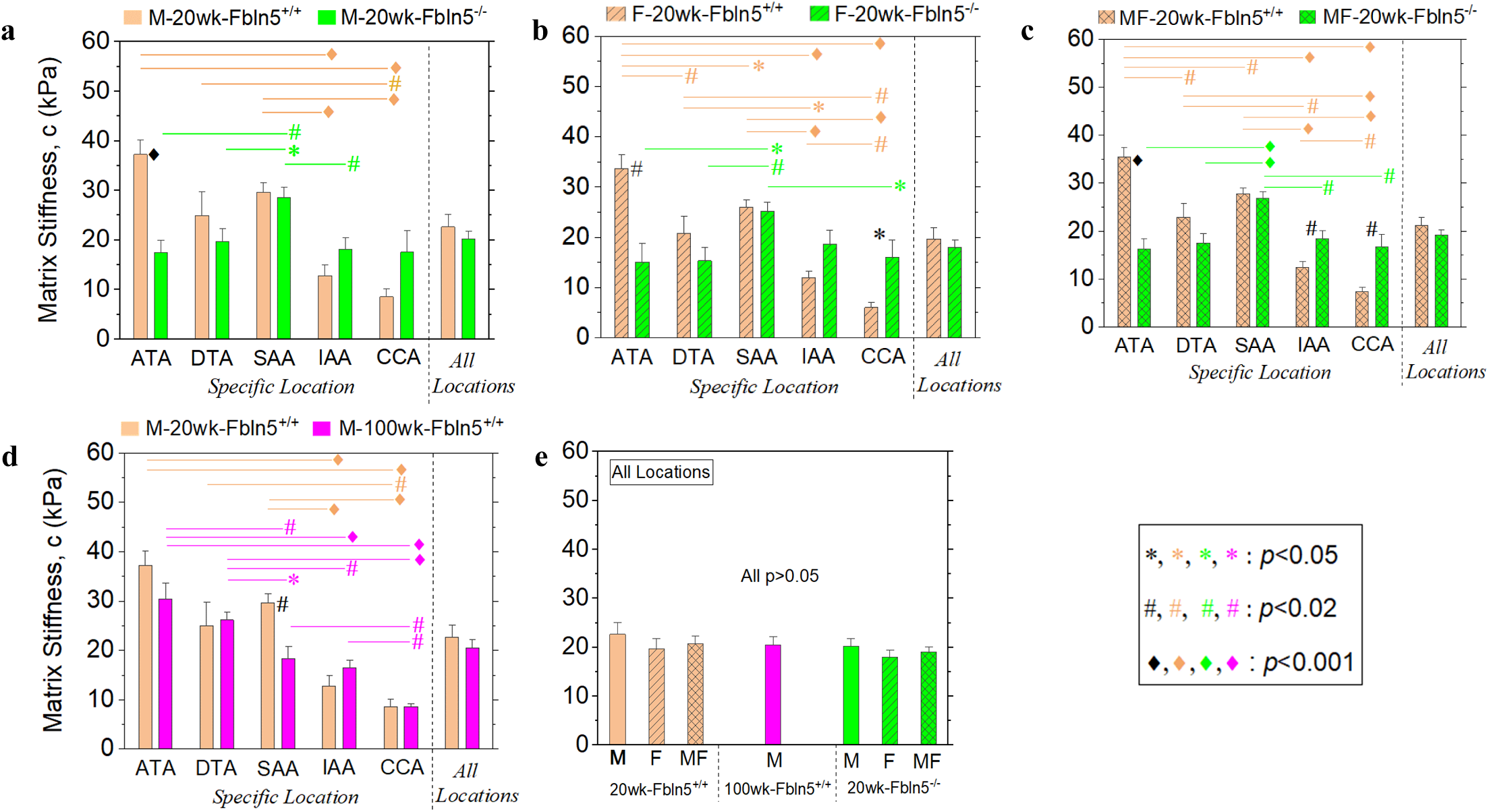
Mean and standard error of arterial matrix stiffness of male (M) and female (F) mice at 20 weeks (20wk) and 100 weeks (100wk) old, for wild-type (Fbln5^+/+^) and knock-out type (Fbln5^-/-^). ATA: ascending thoracic aorta, DTA: descending thoracic aorta, SAA: suprarenal abdominal aorta, IAA: infrarenal abdominal aorta, and CCA: common carotid artery. MF: male and female analyzed together. Asterisks 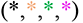: *p*<0.05, Hash signs 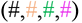: *p*<0.02, and Diamonds 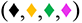: *p*<0.001. Black symbols (*, #, ♦) indicate statistic difference between different genotypes and/or different ages. Colored symbols 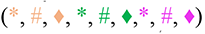 near the right end of a line indicate statistic difference between different locations. Arterial matrix stiffness follows similar patterns in terms of location for both M and F, with either Fbln5^+/+^ or Fbln5^-/-^ (**a**, **b**, **c**). It decreases from ATA to SSA to CCA for Fbln5^+/+^ groups (orange bars in **a**, **b**, **c**), while first increases from ATA to SSA, and then decreases to CCA for Fbln5^-/-^ groups (green bars in **a**, **b**, **c**). The Fbln5 knockout decreases the matrix stiffness at ATA (**a**, **b**, **c**), while increases the matrix stiffness at IAA and CCA (**c**). The aging effect has not much effect on the matrix stiffness at most locations (only reduces the value at SAA, in **d**).

Aging caused a significant decrease in the circumferential fiber component (Fig. 4d, locations of DTA, SAA, and CCA). The circumferential fiber component was not different between wild type and Fbln5^-/-^ arteries at most locations (only reducing the value at SAA of male, Fig. 4a). Aging effect and Fbln5 knockout had little influence on the fiber initial stiffness (Fig. 5), which indicates that the microstructure changes of elastic fibers (Fig. 2) may not affect the mechanical response at small deformation (e.g., <5% in strain).

**Fig 4.**
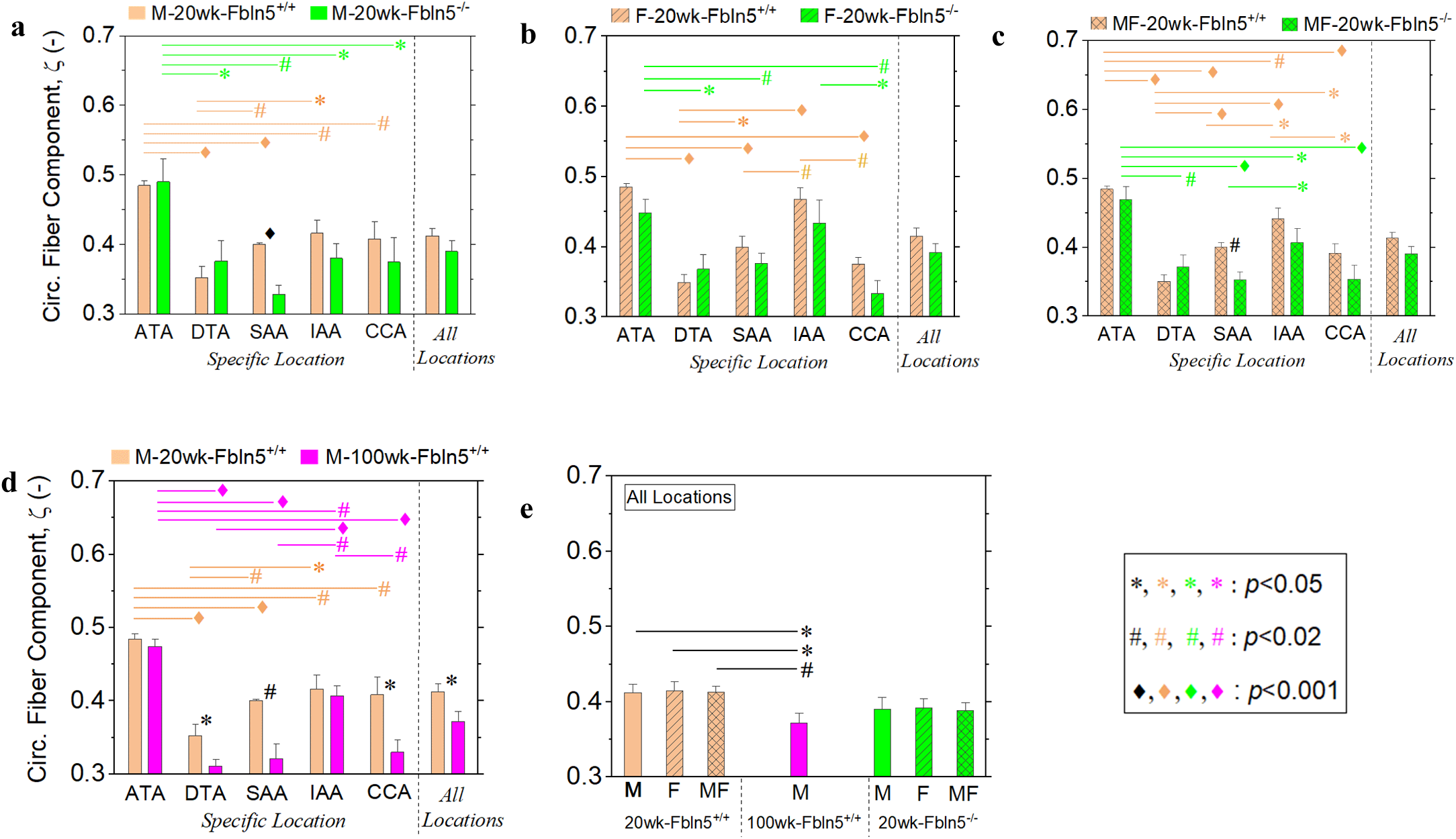
Mean and standard error of circumferential (Circ.) fiber component of male (M) and female (F) mice at 20 weeks (20wk) and 100 weeks (100wk) old, for wild-type (Fbln5) and knock-out type (Fbln5). ATA: ascending thoracic aorta, DTA: descending thoracic aorta, SAA: suprarenal abdominal aorta, IAA: infrarenal abdominal aorta, and CCA: common carotid artery. MF: male and female analyzed together. Asterisks 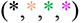: *p*<0.05, Hash signs 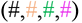: *p*<0.02, and Diamonds 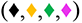: *p*<0.001. Black symbols (*, #, ♦) indicate statistic difference between different genotypes and/or different ages. Colored symbols 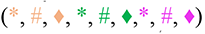 near the right end of a line indicate statistic difference between different locations. Circumferential fiber component follows similar patterns in terms of location for both M and F, with either Fbln5^+/+^ or Fbln5^-/-^ (**a**, **b**, **c**). The aging effect significantly decreases circumferential fiber component (locations of DTA, SAA, and CCA in **d**), while the Fbln5 knockout has not much effect at most locations (only reduce the value at SAA of male, in **a**).

**Fig 5.**
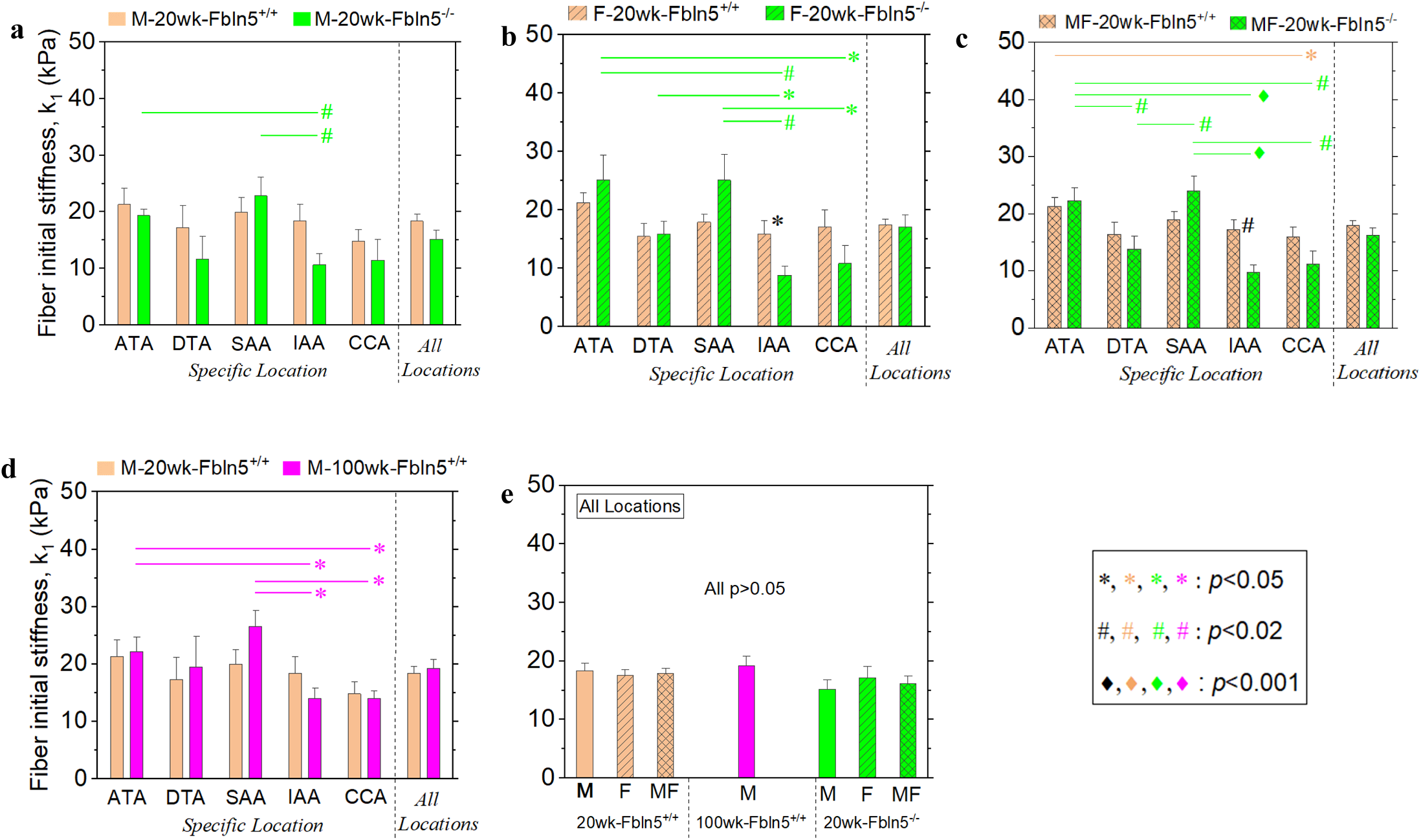
Mean and standard error of arterial fiber initial stiffness of male (M) and female (F) mice at 20 weeks (20wk) and 100 weeks (100wk) old, for wild-type (Fbln5) and knock-out type (Fbln5). ATA: ascending thoracic aorta, DTA: descending thoracic aorta, SAA: suprarenal abdominal aorta, IAA: infrarenal abdominal aorta, and CCA: common carotid artery. MF: male and female analyzed together. Asterisks 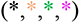: *p*<0.05, Hash signs 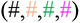: *p*<0.02, and Diamonds 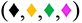: *p*<0.001. Black symbols (*, #, ♦) indicate statistic difference between different genotypes and/or different ages. Colored symbols 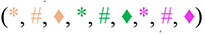 near the right end of a line indicate statistic difference between different locations.

## DISCUSSIONS

This study provided a comprehensive quantitative analysis of the effect of aging, Fbln5 gene, and physiological locations on the arterial stiffness of mice. The results showed that both aging and loss-of-function of fibulin-5 result in statistically significant changes in the arterial stiffening of these arteries, which provides new insights to reduce/treat the adverse stiffening effect of Fbln5 gene mutation and natural aging. For instance, gene corrections/edits may be promising methods to reverse the arterial stiffening and ‘arterial age’ caused by Fbln5 gene mutation. This study offered an accurate characterization of arterial stiffness, and provided guidance on which parameter/property (e.g., *k*_2_) needed to be focused on during the experiments of gene corrections/edits. To our best knowledge, our UFD model^29–31^ applied here is the only existing model that could capture the property/stiffness differences between different groups of the experimental data^14, 15^ discussed in this work. Without the current work, it may be even difficult to identify whether the gene corrections/edits have influences on the arterial stiffness during practice.

It should be noted that the 4-fiber family model^32^ was applied to analyze the experimental data^14, 15^; however, the 4-fiber family model did not capture statistically significant differences with aging or fibulin-5 loss of function. The reason could be the intrinsic inconsistency between the assumption of 4-fiber-family distribution and the real arterial fiber distribution: 1) the number of fiber families in arterial tissue may not be the number four and could even vary at different physiological locations^38^. Similar issues exist in other fiber-family-based models such as the well-known Gasser-Ogden-Holzapfel (GOH) model^23^ which is based on two fiber families; and 2) the 4-fiber-family model assumes all the fibers are distributed at four directions (axial, circumferential and two symmetric diagonal directions with respect to axial), while the fibers could disperse in any direction within the tissue plane^38–41^. In contrast, the UFD model considers the fibers as a unified distribution, which can be applied to arbitrary planar fiber distributions (with any number of fiber families) as long as an in-plane symmetric axis exists. Such consideration is more physically consistent with the real fiber distribution of arterial tissues, because there are two natural symmetries (axial and circumferential directions) in arterial tissues and the majority fibers are distributed within the tissue plane^38^.

In addition, the 4-fiber family model^14, 15, 32^ has 8 material parameters which may result in the overparameterization issue during the fitting, in contrast to the UFD model^29–31^ which has 4 material parameters but still achieves good accuracy (Fig. 1a, b, and Extended Data Fig. 1). If we regard a constitutive model as one kind of transform, different constitutive models will transform the experimental data into different spaces spanned by their corresponding parameters. Due to its physical-consistent assumption, it is more clear to see the experimental data from the space spanned by the parameters of the UFD model, as shown in our current results. Thus, it may be worth using the recently developed UFD model to re-check the huge amount of mechanical testing data in the literature^12, 13, 16, 17, 42–44^, among others.

To our best knowledge, this is also the first study that reports the arterial property/stiffness of male and female mice with the same genotype following the same pattern in terms of locations (Fig. 1 c-e). The material property of the aorta/arteries is highly nonhomogeneous in terms of location (as shown here and by others^21, 22^). It is still a challenging problem to identify the nonhomogeneous *in vivo* properties of arterial/aortic walls, and most existing inverse studies^45, 46^ assumed a homogeneous property of arterial/aortic walls in terms of location. The location pattern of the central arteries found in this study may allow us to estimate the entire property field at different locations based on the property of only one location.

One limitation in this work is that we considered the testing tissue samples as homogeneous, and materially uniform. Thus, the results represented a through-thickness average for the property of the arterial tissue. Moreover, the fitting results here may correspond to a local optimum. Currently, for general nonlinear regression problems with multiple fitting parameters, there is no feasible method that could guarantee a global optimum^47^. The number of local optimum usually increases with increasing number of fitting parameters. Considering that the UFD model^29–31^ reduced the parameter number to only four, in comparison to the previous 4-fiber-family model^32^ with eight parameters, the current fitting results may have a much less possibility for a local optimum. Finally, we only have male mice for the aging group (100 weeks old). Future work may also include female mice at 100 weeks old to investigate the potential aging difference between male and female.

In summary, this study revealed the effect of aging, sex, Fbln5 gene, and physiological locations on the intrinsic arterial stiffening of mice. Both the aging and knockout of Fbln5 gene significantly increase the arterial stiffness, with the latter having a more serious effect. We also showed that 20 weeks mice (equivalent to ∼26 years old human^35^) with Fbln5^-/-^ are older, with respect to ‘arterial age’, than 100 weeks old wild-type (Fbln5^+/+^) mice (equivalent to ∼77 years old human^35^). No statistical difference was found between the male and female groups. In addition, the arterial stiffness of male and female mice follows the same pattern among the five locations (ATA, DTA, SAA, IAA and CCA). The methods offered an accurate characterization of arterial stiffness and the findings provided new insights to reverse/reduce arterial age caused by abnormal Fbln5 gene mutations and natural aging.

## Methods

### Unified-fiber-distribution (UFD) model

We assumed that the arterial tissues are incompressible^23, 24^, and used our recently developed unified-fiber-distribution (UFD) model^29–31^ to characterize the mechanical behavior of the arteries. The fibers in the arterial tissue are usually not well-aligned in one direction, but dispersed in different directions^38–41^. The UFD model considers the fibers in the arterial tissue as a unified distribution, which are more physically consistent with the real fiber distribution of arterial tissues than separating the fiber distribution into several fiber families^23, 24, 32, 33^. When applied to arterial tissues, the strain energy function of the UFD model can be expressed as

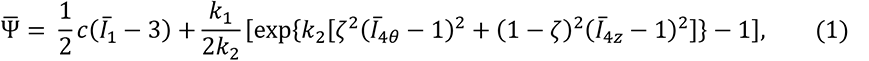

where the first term is the matrix contribution characterized by the neo-Hookean model^48–54^, with the parameter *c* representing the matrix stiffness (see Fig. 6c). The second term in Eq. (1) is the fiber contribution which distinguishes the UFD model from other models. The parameters *k*_1_, *k*_2_ and ζ are fiber related material parameters with *k*_1_ representing the initial stiffness (Fig. 6a), *k*_2_ representing the stiffening effect under stretch (Fig. 6b), and ζ is a scalar characterizing the fiber distribution, representing the average circumferential fiber component. ζ also controls the degree of anisotropy between axial and circumferential directions (Fig. 6d). *Ī*_1_ = tr(C̄) is the 1st invariant of the right Cauchy–Green tensor **C̄**. The invariants *Ī*_4θ_ and *Ī*_4z_ represent the deformation along the circumferential and axial directions, respectively. They are given by *Ī*_4θ_ = (**a**_0_*_θ_* ⊗ **a**_0_*_θ_*): **C̄**, and *Ī*_4z_ = (**a**_0_*_z_* ⊗ **a**_0_*_z_*): **C̄**, where {**a**_0_*_θ_*, **a**_0_*_z_*} are the unit vectors in the circumferential and axial directions, respectively.

**Fig 6.**
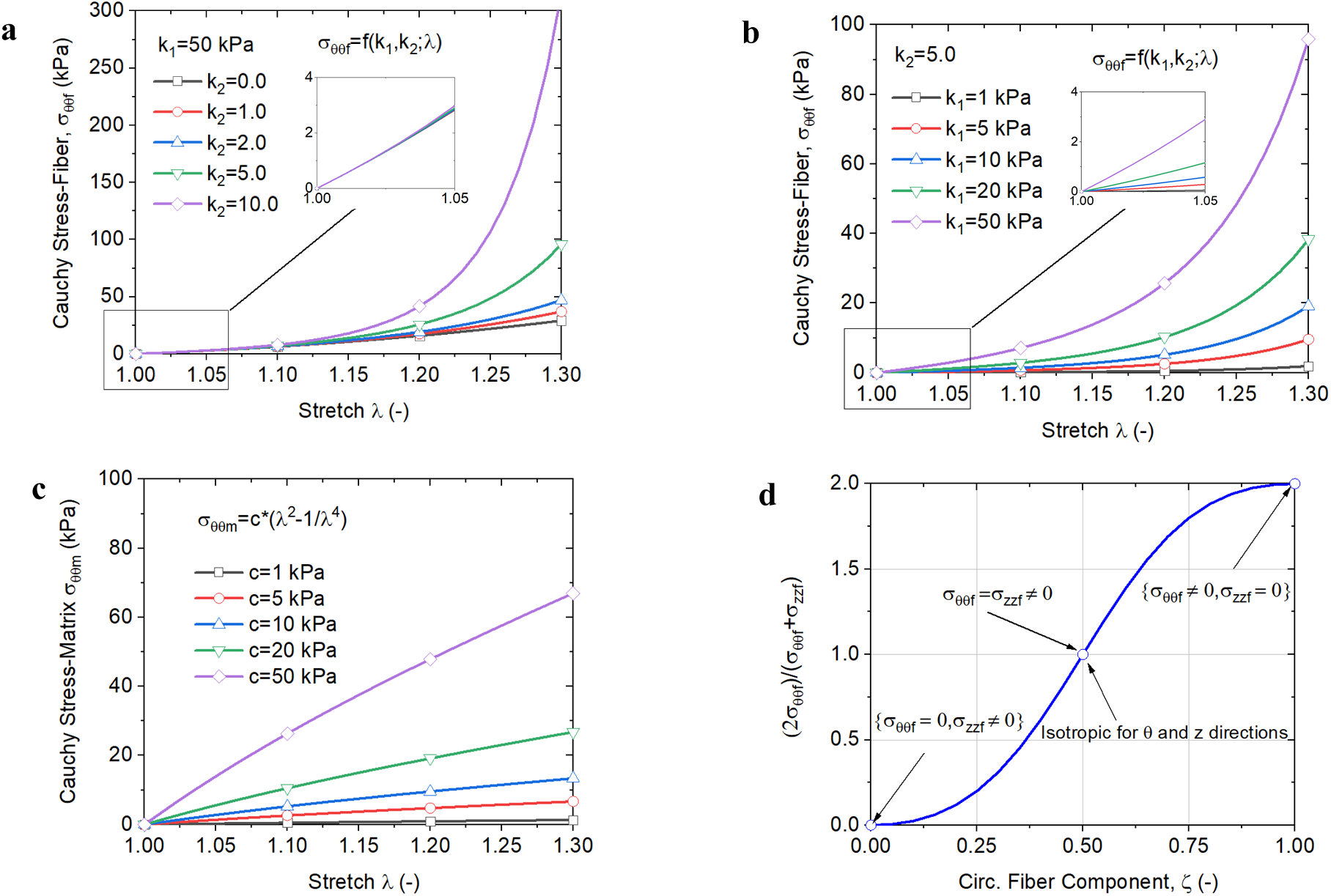
Cauchy stress under equal biaxial tension (*λ*_θ_ = *λ*_z_ = *λ*) in axial (z) and circumferential (*θ*) directions. **a-b**: Fiber contribution to the stress-stretch relation. **a**: fixed *k*_1_ with changed *k*_2_, the initial slope (initial modulus) of the curve does not change, while the stiffening effect at large deformation increases with increasing *k*_2_; **b**: changed *k*_2_ with fixed *k*_1_, the initial slope of the curve increases proportionally with increasing *k*_1_, while the stiffening effect (ratio between slope at large deformation and its initial value) does not change. **c**: Matrix contribution to the stress-stretch relation: there is little stiffening effect at large deformation from the matrix contribution. (**a-c)** indicate the parameter *c* controls the stiffness of the matrix, *k*_1_ controls the initial stiffness of the fiber, and *k*_2_ controls the stiffening at large deformation of the entire tissue (matrix has little stiffening effect). In **a-b**, the parameter ζ = 0.5. **d**: ratio of (2*σ_θθf_*)/*(σ_θθf_* + *σ_zzf_*) on the parameter ζ ∈ [0,1]. ζ represents the average fiber component in the circumferential (*θ*) direction. (1 – ζ) represents the average fiber component in the axial (*z*) direction. ζ = 0 represents the distribution of all the fibers along the axial direction (*σ_θθf_* = 0, *σ_zzf_* ≠ 0), ζ = 1 corresponds to the distribution of all the fibers along the circumferential direction (*σ_θθf_* = 0, *σ_zzf_* ≠ 0), and ζ = 0.5 corresponds to the same fiber component along the axial and circumferential directions (*σ_θθf_* = *σ_zzf_* ≠ 0, isotropic for axial and circumferential). Thus, ζ controls the degree of anisotropy. The explicit form of *σ_θθf_* and *σ_zzf_* are given by Eq. (SE10) in the Supplementary Information.

It is usually considered that the fibers in aortic tissues can support tension loadings but cannot support compression loadings^23, 24^. A more general form of the strain energy function in Equation (1) may be written as

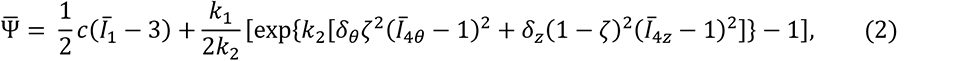

where the role of *δ_θ_* and *δ_z_* is to exclude the contribution of the compressed fibers, given by the Heaviside step function

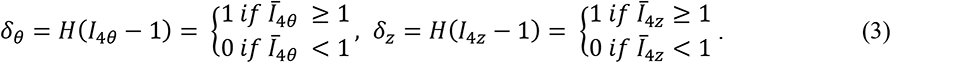

### Biaxial testing data

We have a total of 128 biaxial testing samples of mouse central arteries, which were obtained from Ferruzzi et al.^14, 15^. The samples were classified into three data pairs (**Table 1**):

1. Pair-1 includes two groups with (Fbln5^+/+^) and without (Fbln5^-/-^) Fbln5 gene for male (M) mice at 20 weeks old (20wk);
2. Pair-2 includes two groups of Fbln5^+/+^ and Fbln5^-/-^ for female (F) mice at 20wk;
3. Pair-3 includes two groups at 20 and 100 weeks old for M mice of Fbln5^+/+^.

**Table 1:** Three pairs of biaxial testing data of mouse arteries. M: male, F: female, 20wk/100wk: 20/100 weeks old, Fbln5^+/+^: wild-type with fibulin-5 gene, Fbln5^-/-^: knockout type without fibulin-5 gene. ATA: ascending thoracic aorta, DTA: descending thoracic aorta, SAA: suprarenal abdominal aorta, IAA: infrarenal abdominal aorta, and CCA: common carotid artery. Note that Pair-1 and Pair-3 contain a same group type (M-20wk-Fbln5^+/+^).

| Data Pair | Group type | Specific Location |  |  |  |  |
| --- | --- | --- | --- | --- | --- | --- |
|  |  | ATA | DTA | SAA | IAA | CCA |
| Pair-1 | $M-20wk-Fbln5^{+/+}$ | n=5 | n=5 | n=5 | n=5 | n=5 |
| | $M-20wk-Fbln5^{-/-}$ | n=5 | n=5 | n=5 | n=5 | n=5 |
| Pair-2 | $F-20wk-Fbln5^{+/+}$ | n=5 | n=5 | n=5 | n=5 | n=5 |
| | $F-20wk-Fbln5^{-/-}$ | n=5 | n=5 | n=5 | n=5 | n=5 |
| Pair-3 | $M-20wk-Fbln5^{+/+}$ | n=5 | n=5 | n=5 | n=5 | n=5 |
| | $M-100wk-Fbln5^{+/+}$ | n=6 | n=6 | n=5 | n=6 | n=5 |

Note that Pair-1 and Pair-3 contain a common group (M-20wk-Fbln5^+/+^).

Each group in **Table 1** contains five sub-groups of five different locations including ascending thoracic aorta (ATA), descending thoracic aorta (DTA), suprarenal abdominal aorta (SAA), infrarenal abdominal aorta (IAA), and common carotid artery (CCA). Each sub-group has 5 or 6 samples (**Table 1**) and, for each sample, the biaxial testing data contain a total of seven protocols including three protocols of Pressure-Diameter testing at different fixed axial stretches (Fig. 1a), and four protocols of Force-Length testing at different fixed pressures (Fig. 1b).

### Parameter estimation

The values of *c*, *k*_1_, *k*_2_ and ζ for each sample were obtained by fitting the seven protocols of the biaxial testing data, with minimizing the following error function^12, 13^

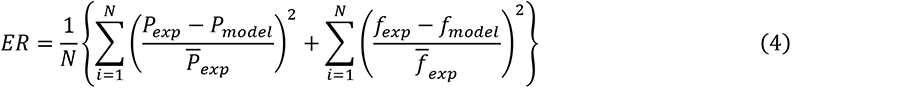

where *N* is the total number of the data point of the seven protocols of one sample. *P_exp_* and *f_exp_* are the measured pressure and non-reduced axial force, respectively, of each data point from experiments. *P*_model_ and *f_model_* are the theoretical pressure and non-reduced axial force, respectively, calculated based on the UFD model (see Supplementary Information). *P̄_exp_* and *f̄_exp_* are the mean pressure and mean non-reduced axial force of the *N* data points, respectively. The *lsqnonlin* function for nonlinear least-squares fitting in MATLAB 2018b (Mathworks Inc., Natick, MA) was applied for the parameter fitting procedure.

### Histology of elastic fibers

We fixed the samples overnight in 4% formalin while unloaded, then stored in 70% ethanol at 4 °C until embedding in paraffin and sectioning at 5 *μ*m. To identify elastic fibers, sections were stained with Verhoeff van Gieson (VVG) and imaged using an Olympus BX/51 microscope with an either 20× or 40× objective. Complete cross sectional views of the arterial wall were obtained by stitching individual images acquired via an Olympus DP70 digital camera and software (CELLSENS DIMENSION).

### Statistical analysis

Both parametric test (Two-Sample t-test), and nonparametric test (Wilcoxon rank sum test) were used to compare the parameters from different groups, with the null hypothesis that the parameters come from independent random populations. The analyses were performed with the functions of *ttest2* (Two-Sample t-test) and *ranksum* (Wilcoxon rank sum test) in MATLAB 2018b (Mathworks Inc., Natick, MA), A *p*-value less than 0.05 was considered to be statistically significant, with three categories marked in Figs. 1-5 and Extended Data Figs. 2-5: i) *p*<0.05, marked as asterisks; ii) *p*<0.02, as hash signs; and iii) *p*<0.001, as diamonds.

## Acknowledgments

The authors would like to thank Dr. Jay D. Humphrey for his kind support. H.D. would like to thank Dr. Xiaoyue Ni, Dr. Enzheng Shi, Dr. Max Lau, Mr. Wei Wang, and Mr. Jiahe Huang for their valuable discussions. H.D. also thanks Mr. Samuel Dembowitz, Ms. Zhenru Li, and Ms. Yumei Shang for their assistance on data analyses. This study is supported by NIH (R01HL155537 and R01HL143348).

## Author contributions

H.D.: Conceptualization; Data curation; Formal analysis; Investigation; Methodology; Software; Validation; Writing—original draft. J.F.: Conceptualization; Data curation; Formal analysis; Investigation; Methodology; Writing—review & editing. M.L.: Investigation; Software; Writing—review & editing. L.P.B: Conceptualization; Writing—review & editing. B.G.L: Conceptualization; Funding acquisition; Writing—review & editing. R.L.G.: Conceptualization; Funding acquisition; Investigation; Project administration; Resources; Supervision; Writing— review & editing.

## Competing interests

The authors declare that they have no known competing financial interests or personal relationships that could have appeared to influence the work reported in this paper.

---

**Extended Data Fig 1.**
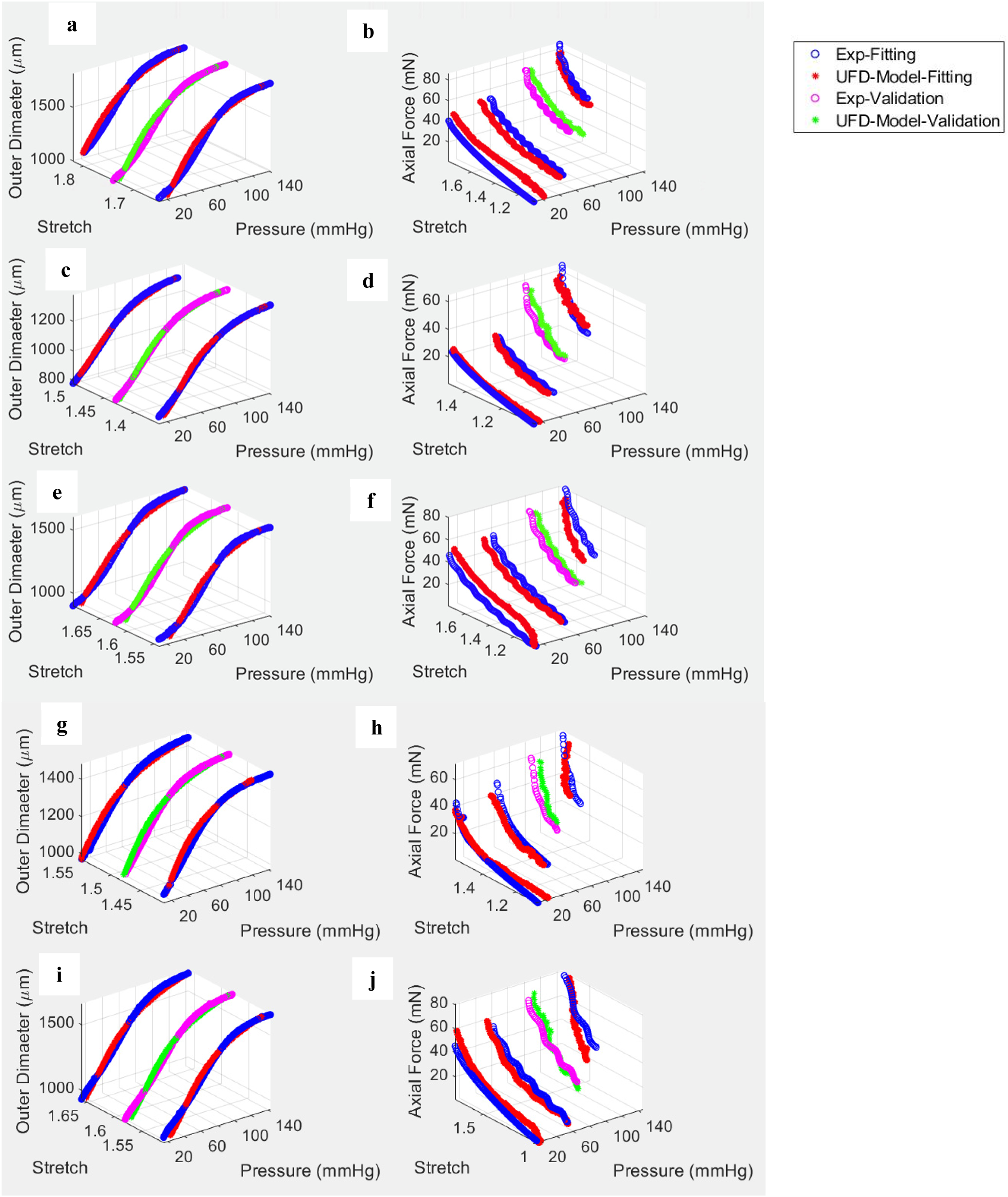
Five representative samples of accuracy validation of the UFD model^6–8^, by predicting the results of the two protocols (Exp-Validation) based on fitted the parameters from the other five (Exp-Fitting) of the seven protocols (two figures in a row). The predicted results also agree very well with the experiments.

**Extended Data Fig 2.**
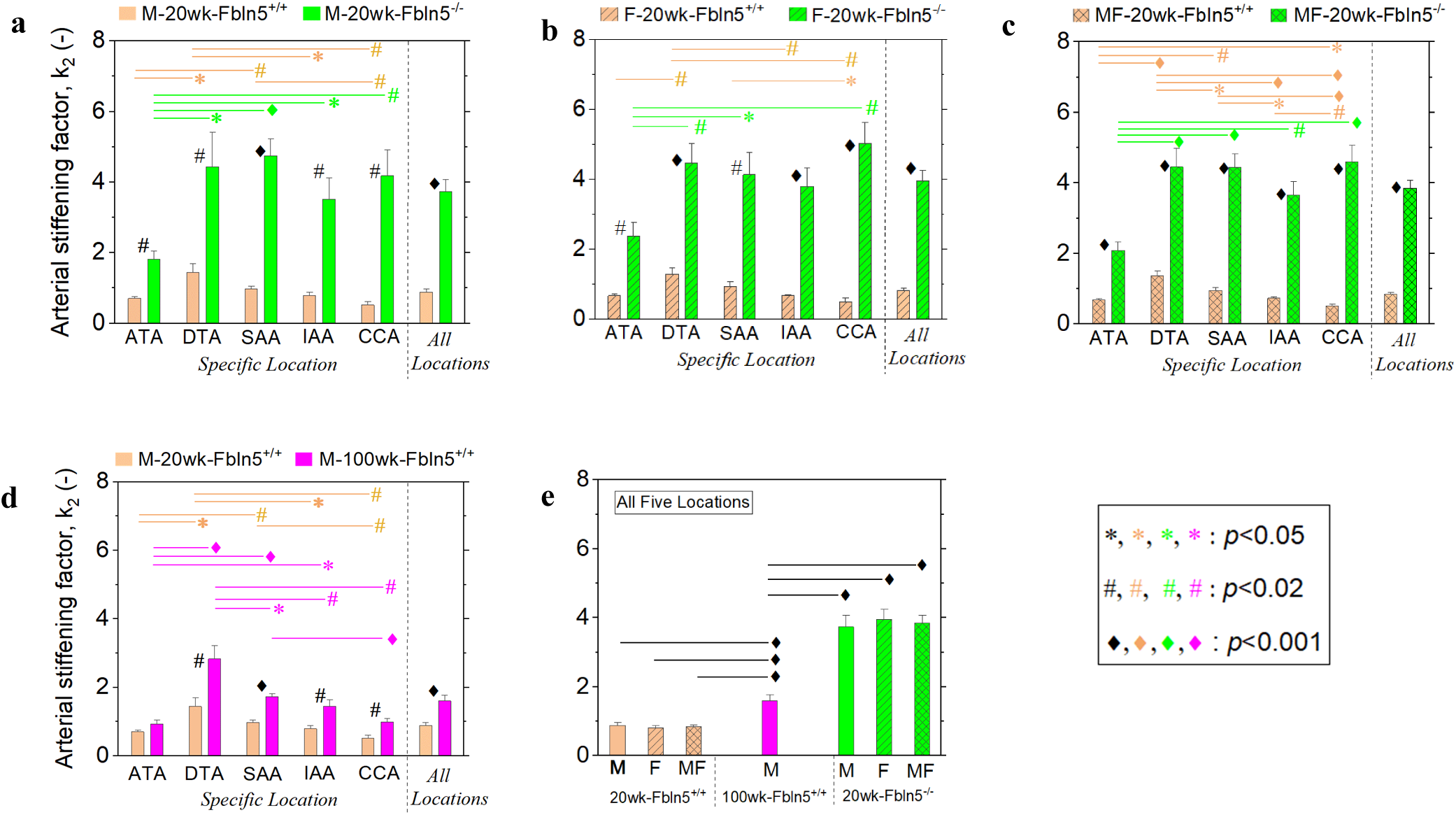
Mean and Standard error of arterial stiffening factor (*k*_2_) of male (M) and female (F) mice at 20 weeks (20wk) and 100 weeks (100wk) old, for wild-type (Fbln5^+/+^) and knock-out type (Fbln5^-/-^). ATA: ascending thoracic aorta, DTA: descending thoracic aorta, SAA: suprarenal abdominal aorta, IAA: infrarenal abdominal aorta, and CCA: common carotid artery. MF: male and female analyzed together. *p-*vlaues based on Wilcoxon rank sum test. Asterisks 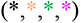: *p*<0.05, Hash signs 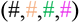: *p*<0.02, and Diamonds 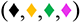: *p*<0.001. Black symbols (*, #, ♦) indicate statistic difference between different genotypes and/or different ages. Colored symbols 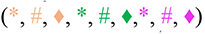 near the right end of a line indicate statistic difference between different locations. Arterial stiffening with Fbln5^-/-^ is significantly larger than that with Fbln5^+/+^, for both M and F, at each of the five locations (**a**, **b, c**). The aging effect also significantly increases the arterial stiffening (**d**). Arterial stiffening at 20 weeks old with Fbln5^-/-^ is even larger than that at 100 weeks old with Fbln5^+/+^ (**e**), indicating that the 20 weeks old mice (equivalent to ∼26 years old human^11^) of Fbln5^-/-^ have a higher ‘vascular age’ than 100 weeks old wild-type mice (equivalent to ∼77 years old human^11^) of Fbln5^+/+^. Arterial stiffening factor follows the same pattern in terms of locations for M (**a**) and F (**b**), with either Fbln5^+/+^ or Fbln5^-/-^. For Fbln5^+/+^ groups (orange bars in **a**, **b, c**), arterial stiffening factor increases from ATA to DTA, and then decreases to IAA to CCA; For Fbln5^-/-^ groups (green bars in **a**, **b, c**), the arterial stiffening factor increases from ATA to DTA, and then keeps about the same value among DTA, SAA, IAA, and CCA.

**Extended Data Fig 3.**
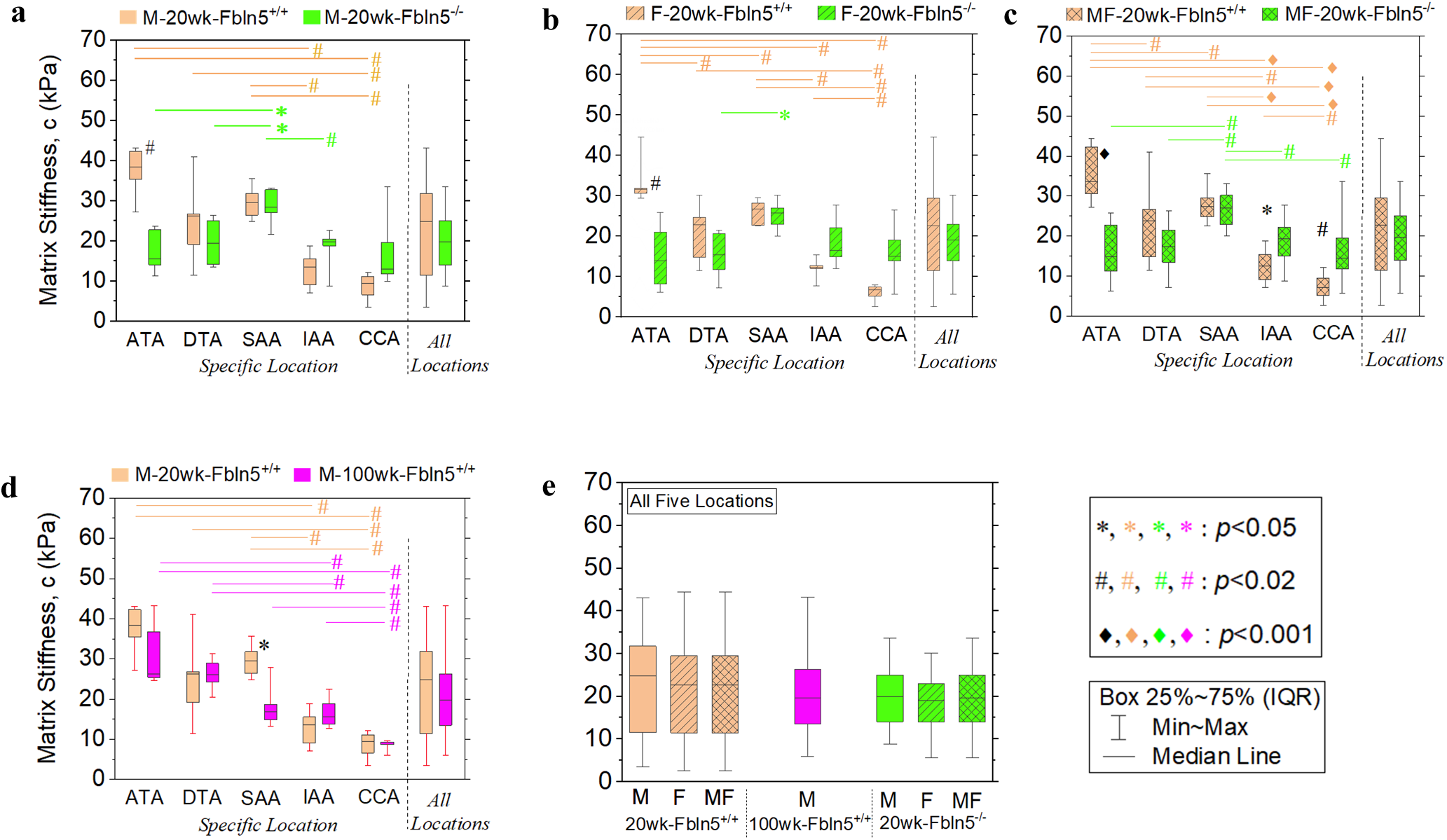
Arterial matrix stiffness of male (M) and female (F) mice at 20 weeks (20wk) and 100 weeks (100wk) old, for wild-type (Fbln5^+/+^) and knock-out type (Fbln5^-/-^). ATA: ascending thoracic aorta, DTA: descending thoracic aorta, SAA: suprarenal abdominal aorta, IAA: infrarenal abdominal aorta, and CCA: common carotid artery. MF: male and female analyzed together. *p-*vlaues based on Wilcoxon rank sum test. Asterisks 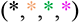: *p*<0.05, Hash signs 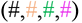: *p*<0.02, and Diamonds 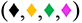: *p*<0.001. Black symbols (*, #, ♦) indicate statistic difference between different genotypes and/or different ages. Colored symbols 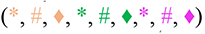 near the right end of a line indicate statistic difference between different locations. Arterial matrix stiffness follows similar patterns in terms of location for both M and F, with either Fbln5^+/+^ or Fbln5^-/-^ (**a**, **b**, **c**). It decreases from ATA to SSA to CCA for Fbln5^+/+^ groups (orange bars in **a**, **b**, **c**), while first increases from ATA to SSA, and then decreases to CCA for Fbln5^-/-^ groups (green bars in **a**, **b**, **c**). The Fbln5 knockout decreases the matrix stiffness at ATA (**a**, **b**, **c**), while increases the matrix stiffness at IAA and CCA (**c**). The aging effect has not much effect on the matrix stiffness at most locations (only reduces the value at SAA, in **d**).

**Extended Data Fig 4.**
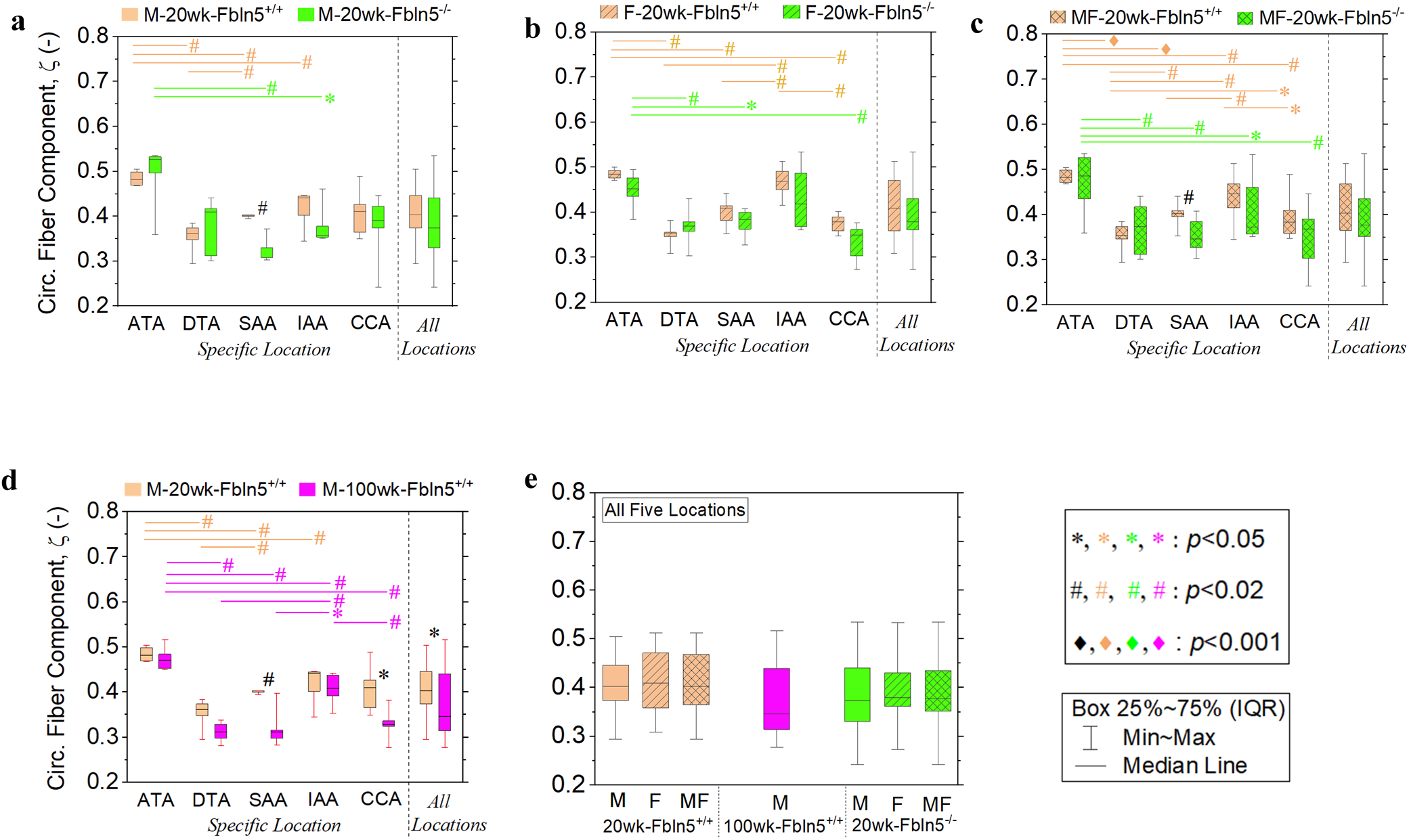
Circumferential (Circ.) fiber component of male (M) and female (F) mice at 20 weeks (20wk) and 100 weeks (100wk) old, for wild-type (Fbln5^+/+^) and knock-out type (Fbln5^-/-^). ATA: ascending thoracic aorta, DTA: descending thoracic aorta, SAA: suprarenal abdominal aorta, IAA: infrarenal abdominal aorta, and CCA: common carotid artery. MF: male and female analyzed together. *p-*vlaues based on Wilcoxon rank sum test. Asterisks 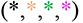: *p*<0.05, Hash signs 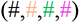: *p*<0.02, and Diamonds 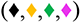: *p*<0.001. Black symbols (*, #, ♦) indicate statistic difference between different genotypes and/or different ages. Colored symbols 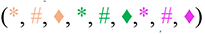 near the right end of a line indicate statistic difference between different locations. Circumferential fiber component follows similar patterns in terms of location for both M and F, with either Fbln5^+/+^ or Fbln5^-/-^ (**a**, **b**, **c**). The aging effect significantly decreases circumferential fiber component (locations of SAA and CCA in **d**), while the Fbln5 knockout has not much effect at most locations (only reduce the value at SAA of male, in **a**).

**Extended Data Fig 5.**
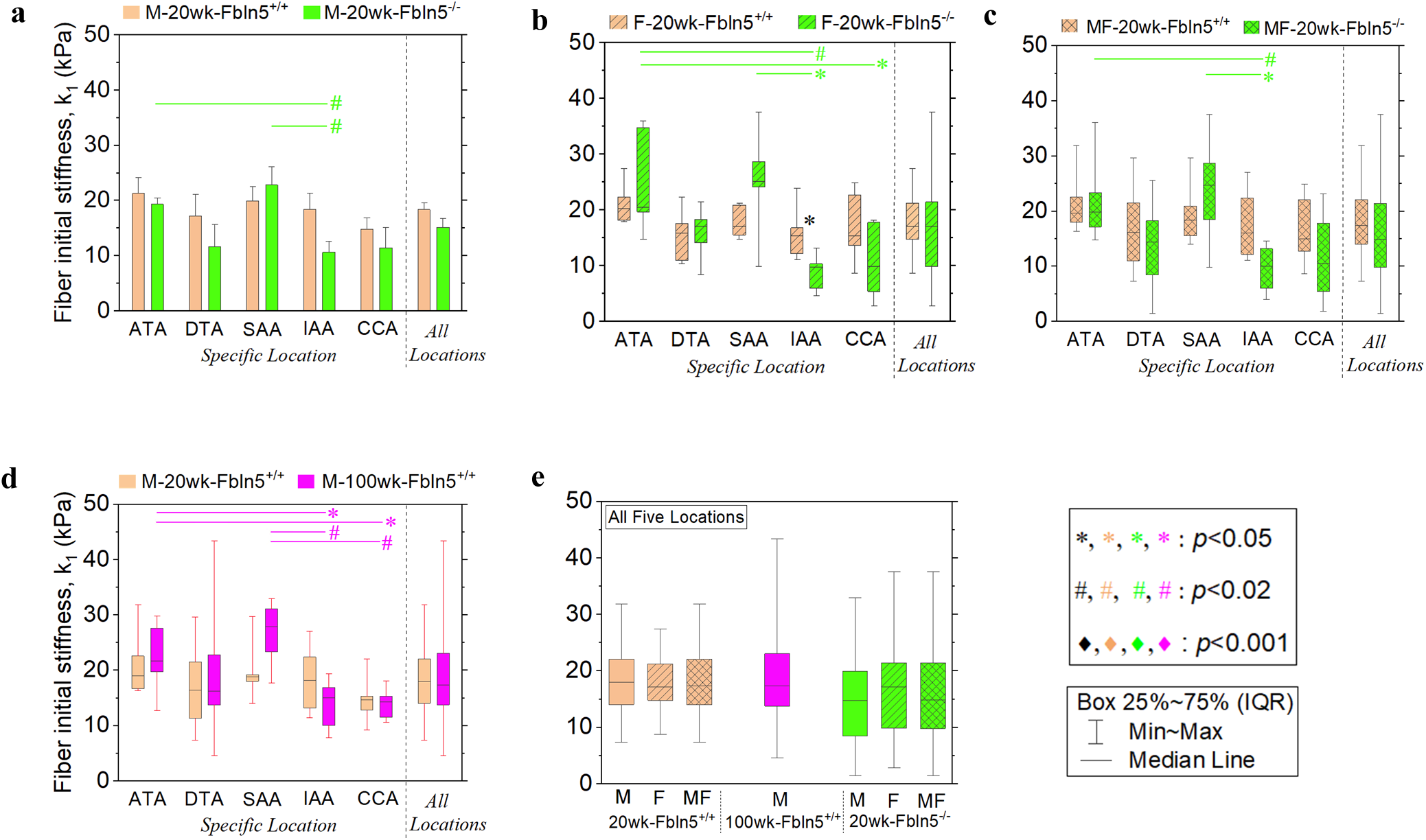
Arterial fiber initial stiffness of male (M) and female (F) mice at 20 weeks (20wk) and 100 weeks (100wk) old, for wild-type (Fbln5^+/+^) and knock-out type (Fbln5^-/-^). ATA: ascending thoracic aorta, DTA: descending thoracic aorta, SAA: suprarenal abdominal aorta, IAA: infrarenal abdominal aorta, and CCA: common carotid artery. MF: male and female analyzed together. *p-*vlaues based on Wilcoxon rank sum test. Asterisks 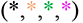: *p*<0.05, Hash signs 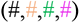: *p*<0.02, and Diamonds 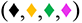: *p*<0.001. Black symbols (*, #, ♦) indicate statistic difference between different genotypes and/or different ages. Colored symbols 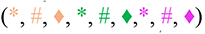 near the right end of a line indicate statistic difference between different locations.

**Extended Data Table 1:**
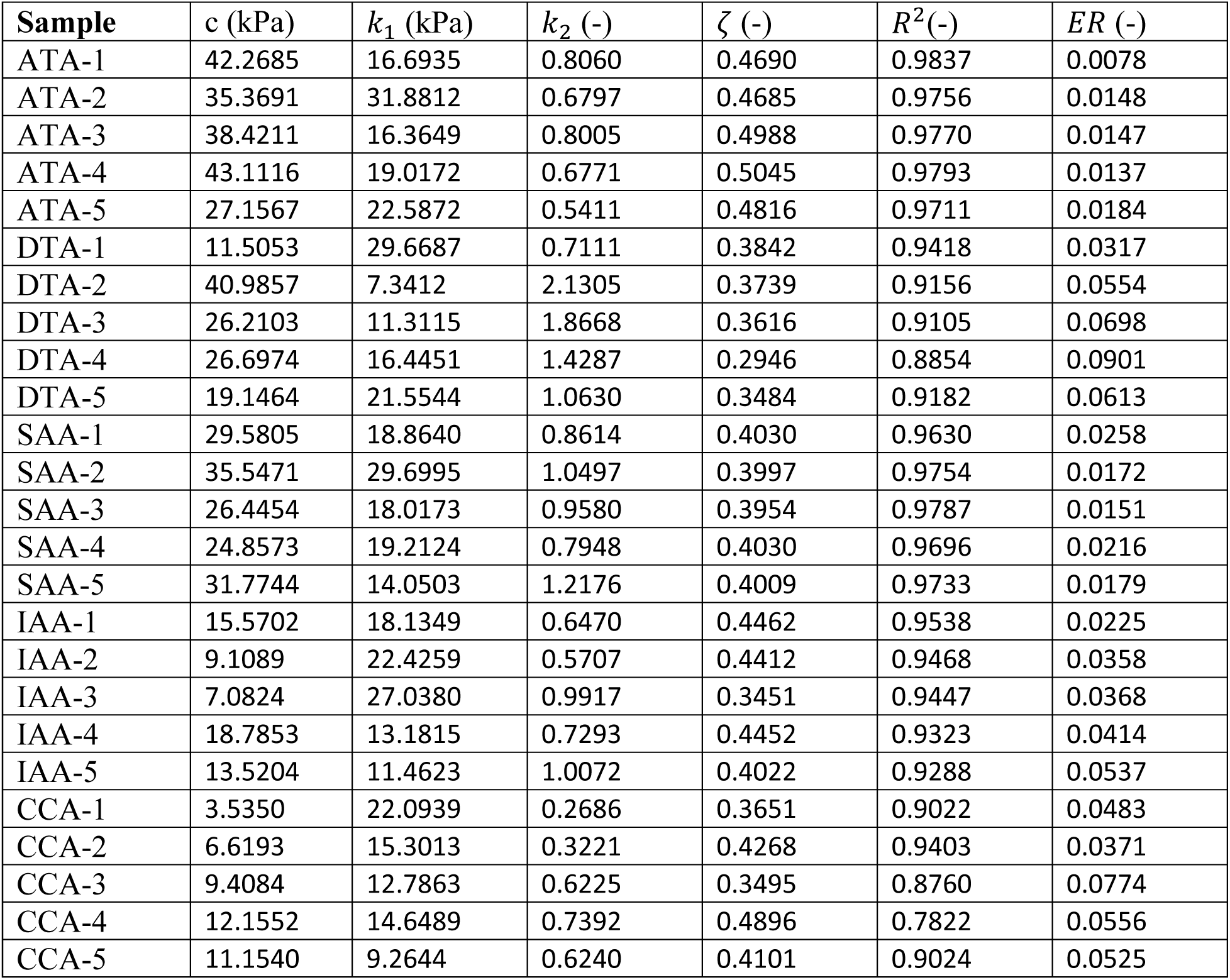
Estimated material parameters *c*, *k*_1_, *k*_2_ and ζ, as well as *R*^2^ and the relative error (*ER*) of each arterial sample of male mice with Fbln5^+/+^ at 20 weeks old (M-20wk- Fbln5^+/+^). ATA: ascending thoracic aorta, DTA: descending thoracic aorta, SAA: suprarenal abdominal aorta, IAA: infrarenal abdominal aorta, and CCA: common carotid artery.

**Extended Data Table 2:** Estimated material parameters *c*, *k*_1_, *k*_2_ and ζ, as well as *R*^2^ and the relative error (*ER*) of each arterial sample of male mice with Fbln5^-/-^ at 20 weeks old (M-20wk- Fbln5^-/-^). ATA: ascending thoracic aorta, DTA: descending thoracic aorta, SAA: suprarenal abdominal aorta, IAA: infrarenal abdominal aorta, and CCA: common carotid artery.

| Sample | $c$ (kPa) | $k_1$ (kPa) | $k_2$ (-) | $\zeta$ (-) | $R^2$(-) | $ER$ (-) |
| --- | --- | --- | --- | --- | --- | --- |
| ATA-1 | 23.6416 | 16.9852 | 2.3155 | 0.4964 | 0.9536 | 0.0351 |
| ATA-2 | 22.7839 | 23.3687 | 2.4394 | 0.5264 | 0.9630 | 0.0269 |
| ATA-3 | 11.2414 | 17.1698 | 1.3674 | 0.5325 | 0.9666 | 0.0232 |
| ATA-4 | 14.0419 | 19.9301 | 1.3376 | 0.5349 | 0.9725 | 0.0229 |
| ATA-5 | 15.5156 | 19.2988 | 1.5781 | 0.3597 | 0.9794 | 0.0117 |
| DTA-1 | 25.0071 | 1.4528 | 7.3494 | 0.3120 | 0.7722 | 0.2595 |
| DTA-2 | 14.1115 | 25.5555 | 2.1715 | 0.4405 | 0.9586 | 0.0274 |
| DTA-3 | 13.4056 | 14.7277 | 2.5075 | 0.4101 | 0.9291 | 0.0678 |
| DTA-4 | 19.4056 | 8.4556 | 4.4192 | 0.3005 | 0.9129 | 0.1035 |
| DTA-5 | 26.3533 | 8.0219 | 5.7447 | 0.4173 | 0.9291 | 0.0708 |
| SAA-1 | 32.7338 | 12.9782 | 4.6383 | 0.3305 | 0.9354 | 0.0648 |
| SAA-2 | 27.0564 | 25.6344 | 5.5244 | 0.3299 | 0.9485 | 0.0495 |
| SAA-3 | 21.7045 | 32.9844 | 3.1102 | 0.3721 | 0.9280 | 0.0686 |
| SAA-4 | 33.1363 | 24.0432 | 5.9129 | 0.3029 | 0.9445 | 0.0530 |
| SAA-5 | 28.4102 | 18.5029 | 4.5349 | 0.3077 | 0.9463 | 0.0544 |
| IAA-1 | 22.7075 | 4.0301 | 5.0852 | 0.3779 | 0.9018 | 0.1141 |
| IAA-2 | 19.8432 | 8.2528 | 2.8209 | 0.3515 | 0.9404 | 0.0565 |
| IAA-3 | 8.8008 | 14.1256 | 1.6152 | 0.3535 | 0.9225 | 0.0637 |
| IAA-4 | 18.7615 | 14.6063 | 3.8683 | 0.3572 | 0.8997 | 0.1012 |
| IAA-5 | 20.4644 | 12.3490 | 4.2067 | 0.4605 | 0.9156 | 0.0748 |
| CCA-1 | 33.5699 | 6.2379 | 5.4795 | 0.4219 | 0.9228 | 0.0547 |
| CCA-2 | 9.8571 | 23.1180 | 2.3632 | 0.3737 | 0.9260 | 0.0597 |
| CCA-3 | 13.0057 | 14.9910 | 3.3410 | 0.3905 | 0.9375 | 0.0458 |
| CCA-4 | 11.7474 | 11.1070 | 3.4140 | 0.4461 | 0.9189 | 0.0733 |
| CCA-5 | 19.5067 | 1.8917 | 6.3037 | 0.2421 | 0.8995 | 0.1031 |

**Extended Data Table 3:** Estimated material parameters *c*, *k*_1_, *k*_2_ and ζ, as well as *R*^2^ and the relative error (*ER*) of each arterial sample of female mice with Fbln5^+/+^ at 20 weeks old (F- 20wk-Fbln5^+/+^). ATA: ascending thoracic aorta, DTA: descending thoracic aorta, SAA: suprarenal abdominal aorta, IAA: infrarenal abdominal aorta, and CCA: common carotid artery.

| Sample | $c$ (kPa) | $k_1$ (kPa) | $k_2$ (-) | $\zeta$ (-) | $R^2$(-) | $ER$ (-) |
| --- | --- | --- | --- | --- | --- | --- |
| ATA-1 | 30.6337 | 20.3042 | 0.4940 | 0.4929 | 0.9675 | 0.0198 |
| ATA-2 | 44.4622 | 27.4242 | 0.7226 | 0.4997 | 0.9809 | 0.0128 |
| ATA-3 | 29.4453 | 17.9630 | 0.6422 | 0.4844 | 0.9672 | 0.0208 |
| ATA-4 | 31.5703 | 22.3113 | 0.8346 | 0.4759 | 0.9721 | 0.0170 |
| ATA-5 | 31.8812 | 18.1470 | 0.6480 | 0.4711 | 0.9701 | 0.0178 |
| DTA-1 | 30.1933 | 10.4215 | 1.6471 | 0.3531 | 0.9256 | 0.0641 |
| DTA-2 | 14.7832 | 15.8754 | 1.3034 | 0.3079 | 0.9272 | 0.0635 |
| DTA-3 | 11.4289 | 22.3480 | 0.6617 | 0.3459 | 0.9097 | 0.0715 |
| DTA-4 | 22.8112 | 17.6483 | 1.3087 | 0.3552 | 0.9509 | 0.0379 |
| DTA-5 | 24.7021 | 11.0179 | 1.5029 | 0.3816 | 0.9327 | 0.0545 |
| SAA-1 | 22.7361 | 21.2670 | 0.5820 | 0.3523 | 0.9551 | 0.0295 |
| SAA-2 | 29.4834 | 14.8037 | 1.3291 | 0.3821 | 0.9623 | 0.0297 |
| SAA-3 | 22.6013 | 15.5418 | 0.6350 | 0.4411 | 0.9640 | 0.0238 |
| SAA-4 | 28.1950 | 20.9080 | 0.9394 | 0.4092 | 0.9715 | 0.0208 |
| SAA-5 | 26.7278 | 17.1375 | 1.1640 | 0.4140 | 0.9532 | 0.0296 |
| IAA-1 | 15.4319 | 12.2009 | 0.7256 | 0.4912 | 0.9321 | 0.0463 |
| IAA-2 | 7.6521 | 16.8081 | 0.6035 | 0.4498 | 0.9255 | 0.0488 |
| IAA-3 | 12.7107 | 23.9681 | 0.7010 | 0.5127 | 0.9280 | 0.0393 |
| IAA-4 | 11.9905 | 15.3334 | 0.6554 | 0.4153 | 0.9261 | 0.0488 |
| IAA-5 | 12.1536 | 11.1261 | 0.6967 | 0.4686 | 0.8954 | 0.0747 |
| CCA-1 | 6.7283 | 13.7373 | 0.5941 | 0.3584 | 0.9235 | 0.0450 |
| CCA-2 | 2.6095 | 15.4499 | 0.4145 | 0.4021 | 0.8923 | 0.0660 |
| CCA-3 | 5.1644 | 22.6978 | 0.3299 | 0.3472 | 0.9226 | 0.0371 |
| CCA-4 | 7.4453 | 8.7209 | 0.8271 | 0.3887 | 0.8846 | 0.0697 |
| CCA-5 | 8.0059 | 24.9199 | 0.3256 | 0.3785 | 0.8737 | 0.0511 |

**Extended Data Table 4:** Estimated material parameters *c*, *k*_1_, *k*_2_ and ζ, as well as *R*^2^ and the relative error (*ER*) of each arterial sample of female mice with Fbln5^-/-^ at 20 weeks old (F-20wk- Fbln5^-/-^). ATA: ascending thoracic aorta, DTA: descending thoracic aorta, SAA: suprarenal abdominal aorta, IAA: infrarenal abdominal aorta, and CCA: common carotid artery.

| Sample | $c$ (kPa) | $k_1$ (kPa) | $k_2$ (-) | $\zeta$ (-) | $R^2$(-) | $ER$ (-) |
| --- | --- | --- | --- | --- | --- | --- |
| ATA-1 | 21.0404 | 34.7427 | 3.6116 | 0.3844 | 0.9840 | 0.0109 |
| ATA-2 | 8.1922 | 36.0619 | 2.0040 | 0.4351 | 0.9673 | 0.0256 |
| ATA-3 | 6.1780 | 14.7956 | 1.3136 | 0.4945 | 0.9785 | 0.0194 |
| ATA-4 | 25.8072 | 20.4723 | 2.8407 | 0.4516 | 0.9825 | 0.0109 |
| ATA-5 | 13.8986 | 19.7276 | 2.1005 | 0.4764 | 0.9674 | 0.0198 |
| DTA-1 | 21.5015 | 17.1649 | 3.8380 | 0.3035 | 0.8561 | 0.1718 |
| DTA-2 | 20.6657 | 21.4295 | 5.5005 | 0.4304 | 0.9213 | 0.0769 |
| DTA-3 | 7.1849 | 14.1908 | 2.9549 | 0.3578 | 0.8666 | 0.1167 |
| DTA-4 | 11.7828 | 18.2800 | 4.0532 | 0.3789 | 0.9035 | 0.1296 |
| DTA-5 | 15.3243 | 8.4894 | 5.9957 | 0.3692 | 0.9137 | 0.0877 |
| SAA-1 | 30.1784 | 9.8660 | 5.3471 | 0.3275 | 0.9225 | 0.0813 |
| SAA-2 | 22.9115 | 24.1991 | 2.9768 | 0.3841 | 0.9432 | 0.0489 |
| SAA-3 | 20.0237 | 28.6851 | 2.3408 | 0.4082 | 0.9119 | 0.0828 |
| SAA-4 | 25.7071 | 37.6030 | 5.6411 | 0.3624 | 0.9393 | 0.0545 |
| SAA-5 | 26.9685 | 25.2101 | 4.3581 | 0.4000 | 0.9467 | 0.0445 |
| IAA-1 | 22.1415 | 9.8188 | 4.1751 | 0.4189 | 0.8981 | 0.1040 |
| IAA-2 | 16.4424 | 13.2475 | 2.5861 | 0.4860 | 0.9005 | 0.0806 |
| IAA-3 | 14.9245 | 10.3362 | 4.8202 | 0.3680 | 0.9217 | 0.0705 |
| IAA-4 | 27.6840 | 4.7137 | 4.9461 | 0.5335 | 0.8870 | 0.0996 |
| IAA-5 | 11.9760 | 6.0012 | 2.4667 | 0.3613 | 0.8955 | 0.1032 |
| CCA-1 | 14.9982 | 9.8893 | 5.0608 | 0.3033 | 0.9171 | 0.0931 |
| CCA-2 | 18.9948 | 2.8493 | 6.4997 | 0.3615 | 0.8895 | 0.1310 |
| CCA-3 | 5.6181 | 18.1472 | 3.2662 | 0.3494 | 0.9021 | 0.0755 |
| CCA-4 | 26.4283 | 17.8144 | 4.2490 | 0.3766 | 0.9416 | 0.0579 |
| CCA-5 | 13.9043 | 5.4498 | 6.0968 | 0.2730 | 0.9046 | 0.0834 |

**Extended Data Table 5:** Estimated material parameters *c*, *k*_1_, *k*_2_ and ζ, as well as *R*^2^ and the relative error (*ER*) of each arterial sample of male mice with Fbln5^+/+^ at 100 weeks old (M- 100wk-Fbln5^+/+^). ATA: ascending thoracic aorta, DTA: descending thoracic aorta, SAA: suprarenal abdominal aorta, IAA: infrarenal abdominal aorta, and CCA: common carotid artery.

| Sample | $c$ (kPa) | $k_1$ (kPa) | $k_2$ (-) | $\zeta$ (-) | $R^2$ (-) | $ER$ (-) |
| --- | --- | --- | --- | --- | --- | --- |
| ATA-1 | 25.8180 | 27.5822 | 0.8766 | 0.4495 | 0.9776 | 0.0112 |
| ATA-2 | 43.2734 | 12.7564 | 1.3945 | 0.5166 | 0.9838 | 0.0107 |
| ATA-3 | 25.4310 | 19.7615 | 0.6790 | 0.4768 | 0.9723 | 0.0137 |
| ATA-4 | 36.8070 | 29.8260 | 1.1868 | 0.4657 | 0.9809 | 0.0097 |
| ATA-5 | 24.5463 | 21.2806 | 0.6761 | 0.4842 | 0.9774 | 0.0147 |
| ATA-6 | 26.7838 | 22.1024 | 0.7543 | 0.4524 | 0.9622 | 0.0162 |
| DTA-1 | 28.8782 | 13.7373 | 2.3585 | 0.3377 | 0.9353 | 0.0659 |
| DTA-2 | 31.2897 | 4.6377 | 4.2058 | 0.2984 | 0.9268 | 0.0835 |
| DTA-3 | 25.0900 | 17.0055 | 1.9165 | 0.2984 | 0.9344 | 0.0684 |
| DTA-4 | 27.0035 | 43.4121 | 3.8790 | 0.2810 | 0.9451 | 0.0445 |
| DTA-5 | 20.4179 | 15.4062 | 2.2293 | 0.3282 | 0.9408 | 0.0467 |
| DTA-6 | 24.1943 | 22.8352 | 2.3911 | 0.3238 | 0.9461 | 0.0481 |
| SAA-1 | 27.8170 | 17.6973 | 1.8681 | 0.3977 | 0.9671 | 0.0239 |
| SAA-2 | 14.8444 | 23.3234 | 1.7923 | 0.3118 | 0.9610 | 0.0310 |
| SAA-3 | 16.7746 | 27.8986 | 1.4881 | 0.3164 | 0.9750 | 0.0194 |
| SAA-4 | 13.2105 | 32.9736 | 1.6002 | 0.2974 | 0.9803 | 0.0116 |
| SAA-5 | 18.6822 | 31.0954 | 1.9065 | 0.2828 | 0.9746 | 0.0209 |
| IAA-1 | 16.6105 | 19.4263 | 1.1564 | 0.3922 | 0.9531 | 0.0302 |
| IAA-2 | 13.7729 | 10.0302 | 1.4152 | 0.3536 | 0.9440 | 0.0469 |
| IAA-3 | 14.4623 | 13.8351 | 1.1985 | 0.4209 | 0.9309 | 0.0557 |
| IAA-4 | 18.9254 | 16.2549 | 1.5797 | 0.3952 | 0.9614 | 0.0244 |
| IAA-5 | 22.4970 | 7.8043 | 2.2931 | 0.4375 | 0.9423 | 0.0366 |
| IAA-6 | 12.6195 | 16.8888 | 1.0754 | 0.4418 | 0.9528 | 0.0304 |
| CCA-1 | 9.3580 | 14.2865 | 0.8032 | 0.3246 | 0.9520 | 0.0329 |
| CCA-2 | 9.3467 | 10.5761 | 1.3626 | 0.2777 | 0.9229 | 0.0614 |
| CCA-3 | 5.9772 | 18.0791 | 0.8502 | 0.3359 | 0.9555 | 0.0247 |
| CCA-4 | 9.5607 | 11.5200 | 0.9688 | 0.3286 | 0.9316 | 0.0415 |
| CCA-5 | 8.6569 | 15.3601 | 0.9384 | 0.3822 | 0.9322 | 0.0441 |

## Notes

### Competing Interest Statement

The authors have declared no competing interest.

## References

1 Arnett, D. K., Evans, G. W. & Riley, W. A. Arterial stiffness: a new cardiovascular risk factor? American journal of epidemiology 140, 669–682 (1994).

2 Vlachopoulos, C., Aznaouridis, K. & Stefanadis, C. Prediction of cardiovascular events and all-cause mortality with arterial stiffness: a systematic review and meta-analysis. Journal of the American College of Cardiology 55, 1318–1327 (2010).

3 Bhuva, A. N. et al. Training for a first-time marathon reverses age-related aortic stiffening. Journal of the American College of Cardiology 75, 60-71 (2020).

4 Thijssen, D. H., Carter, S. E. & Green, D. J. Arterial structure and function in vascular ageing: are you as old as your arteries? The Journal of physiology 594, 2275–2284 (2016).

5 Adji, A., O’rourke, M. F. & Namasivayam, M. Arterial stiffness, its assessment, prognostic value, and implications for treatment. American journal of hypertension 24, 5–17 (2011).

6 Chirinos, J. A., Segers, P., Hughes, T. & Townsend, R. Large-artery stiffness in health and disease: JACC state-of-the-art review. Journal of the American College of Cardiology 74, 1237–1263 (2019).

7 Yanagisawa, H. et al. Fibulin-5 is an elastin-binding protein essential for elastic fibre development in vivo. Nature 415, 168–171 (2002).

8 Nakamura, T. et al. Fibulin-5/DANCE is essential for elastogenesis in vivo. Nature 415, 171–175 (2002).

9 Loeys, B. et al. Homozygosity for a missense mutation in fibulin-5 (FBLN5) results in a severe form of cutis laxa. Human molecular genetics 11, 2113–2118 (2002).

10 Markova, D. et al. Genetic heterogeneity of cutis laxa: a heterozygous tandem duplication within the fibulin-5 (FBLN5) gene. The American Journal of Human Genetics 72, 998–1004 (2003).

11 Tekmenuray-Unal, A. & Durmaz, C. D. FBLN5-Related Cutis Laxa Syndrome: A Case with a Novel Variant and Review of the Literature. Molecular Syndromology, 1–8 (2022).

12 Wan, W., Yanagisawa, H. & Gleason, R. L. Biomechanical and microstructural properties of common carotid arteries from fibulin-5 null mice. Annals of biomedical engineering 38, 3605–3617 (2010).

13 Wan, W. & Gleason Jr, R. L. Dysfunction in elastic fiber formation in fibulin-5 null mice abrogates the evolution in mechanical response of carotid arteries during maturation. American Journal of Physiology-Heart and Circulatory Physiology 304, H674–H686 (2013).

14 Ferruzzi, J., Bersi, M., Uman, S., Yanagisawa, H. & Humphrey, J. Decreased elastic energy storage, not increased material stiffness, characterizes central artery dysfunction in fibulin-5 deficiency independent of sex. Journal of biomechanical engineering 137, 031007 (2015).

15 Ferruzzi, J., Madziva, D., Caulk, A., Tellides, G. & Humphrey, J. Compromised mechanical homeostasis in arterial aging and associated cardiovascular consequences. Biomech Model Mechanobiol 17, 1281–1295 (2018).

16 Martin, C., Sun, W., Pham, T. & Elefteriades, J. Predictive biomechanical analysis of ascending aortic aneurysm rupture potential. Acta biomaterialia 9, 9392–9400 (2013).

17 Pham, T., Martin, C., Elefteriades, J. & Sun, W. Biomechanical characterization of ascending aortic aneurysm with concomitant bicuspid aortic valve and bovine aortic arch. Acta biomaterialia 9, 7927–7936 (2013).

18 Martin, C., Sun, W. & Elefteriades, J. Patient-specific finite element analysis of ascending aorta aneurysms. American Journal of Physiology-Heart and Circulatory Physiology 308, H1306–H1316 (2015).

19 Liu, M. et al. A Novel Anisotropic Failure Criterion With Dispersed Fiber Orientations for Aortic Tissues. Journal of Biomechanical Engineering 142, 111002 (2020).

20 Dong, H. et al. Ultimate tensile strength and biaxial stress–strain responses of aortic tissues—A clinical-engineering correlation. Applications in Engineering Science 10, 100101 (2022).

21 Iliopoulos, D. C. et al. Regional and directional variations in the mechanical properties of ascending thoracic aortic aneurysms. Medical engineering & physics 31, 1–9 (2009).

22 Sokolis, D. P. Layer-Specific Tensile Strength of the Human Aorta: Segmental Variations. Journal of Biomechanical Engineering 145, 064502 (2023).

23 Gasser, T. C., Ogden, R. W. & Holzapfel, G. A. Hyperelastic modelling of arterial layers with distributed collagen fibre orientations. Journal of the royal society interface 3, 15–35 (2006).

24 Holzapfel, G. A., Gasser, T. C. & Ogden, R. W. A new constitutive framework for arterial wall mechanics and a comparative study of material models. Journal of Elasticity the Physical Science of Solids 61, 1–48 (2000).

25 Puck, A. & Schürmann, H. in Failure criteria in fibre-reinforced-polymer composites 832–876 (Elsevier, 2004).

26 Dong, H., Wang, J. & Karihaloo, B. An improved Puck’s failure theory for fibre-reinforced composite laminates including the in situ strength effect. Composites science and technology 98, 86–92 (2014).

27 Dong, H. & Wang, J. A criterion for failure mode prediction of angle-ply composite laminates under in-plane tension. Composite Structures 128, 234–240 (2015).

28 Dong, H., Li, Z., Wang, J. & Karihaloo, B. A new fatigue failure theory for multidirectional fiber-reinforced composite laminates with arbitrary stacking sequence. International Journal of Fatigue 87, 294–300 (2016).

29 Dong, H. & Sun, W. A novel hyperelastic model for biological tissues with planar distributed fibers and a second kind of Poisson effect. Journal of the Mechanics and Physics of Solids 151, 104377 (2021).

30 Dong, H. et al. A novel computational growth framework for biological tissues: Application to growth of aortic root aneurysm repaired by the V-shape surgery. Journal of the Mechanical Behavior of Biomedical Materials, 105081 (2022).

31 Dong, H., Liu, M., Woodall, J., Leshnower, B. & Gleason Jr, R. L. Effect of Nonlinear Hyperelastic Property of Arterial Tissues on the Pulse Wave Velocity based on the Unified-Fiber-Distribution (UFD) Model. bioRxiv, 2022.2009. 2027.509711 (2022).

32 Baek, S., Valentin, A. & Humphrey, J. Biochemomechanics of cerebral vasospasm and its resolution: II. Constitutive relations and model simulations. Annals of biomedical engineering 35, 1498–1509 (2007).

33 Baek, S., Valentín, A. & Humphrey, J. Biochemomechanics of cerebral vasospasm and its resolution: II. Constitutive relations and model simulations. Annals of biomedical engineering 35, 1498–1509 (2007).

34 Henson, G. D., Walker, A. E., Reihl, K. D., Donato, A. J. & Lesniewski, L. A. Dichotomous mechanisms of aortic stiffening in high-fat diet fed young and old B 6 D 2 F 1 mice. Physiological reports 2, e00268 (2014).

35 Dutta, S. & Sengupta, P. Men and mice: relating their ages. Life sciences 152, 244–248 (2016).

36 Hachim, G. & Mdaghri, A. A. Congenital Cutis laxa: Two pediatric cases. World Journal of Advanced Research and Reviews 15, 086–091 (2022).

37 Van Maldergem, L. & Loeys, B. FBLN5-related cutis laxa. GeneReviews®[Internet] (2018).

38 Schriefl, A. J., Zeindlinger, G., Pierce, D. M., Regitnig, P. & Holzapfel, G. A. Determination of the layer-specific distributed collagen fibre orientations in human thoracic and abdominal aortas and common iliac arteries. Journal of the Royal Society Interface 9, 1275–1286 (2012).

39 Canham, P. B., Finlay, H. M., Dixon, J. G., Boughner, D. R. & Chen, A. Measurements from light and polarised light microscopy of human coronary arteries fixed at distending pressure. Cardiovascular research 23, 973–982 (1989).

40 Finlay, H. M., McCullough, L. & Canham, P. B. Three-dimensional collagen organization of human brain arteries at different transmural pressures. Journal of vascular research 32, 301–312 (1995).

41 Finlay, H. M., Whittaker, P. & Canham, P. B. Collagen organization in the branching region of human brain arteries. Stroke 29, 1595–1601 (1998).

42 Lou, X. et al. Biomechanical and Histological Analysis Supports Increased Stiffness and Fibrosis in Chronic versus Acute Aortic Dissection Flaps. Circulation 140, A14347–A14347 (2019).

43 Kim, C. W. et al. Disturbed flow promotes arterial stiffening through thrombospondin-1. Circulation 136, 1217–1232 (2017).

44 Gkousioudi, A. et al. Biomechanical properties of mouse carotid arteries with diet-induced metabolic syndrome and aging. Frontiers in Bioengineering and Biotechnology 10 (2022).

45 Liu, M. et al. Identification of in vivo nonlinear anisotropic mechanical properties of ascending thoracic aortic aneurysm from patient-specific CT scans. Scientific reports 9, 1–13 (2019).

46 Wittek, A. et al. A finite element updating approach for identification of the anisotropic hyperelastic properties of normal and diseased aortic walls from 4D ultrasound strain imaging. Journal of the mechanical behavior of biomedical materials 58, 122–138 (2016).

47 Voit, E. O. A first course in systems biology. (Garland Science, 2017).

48 Holzapfel, G. A. Nonlinear solid mechanics: A continuum approach for engineering science. (John Wiley & Sons, Ltd, 2000).

49 Dong, H., Wang, J. & Rubin, M. A nonlinear cosserat interphase model for residual stresses in an inclusion and the interphase that bonds it to an infinite matrix. International Journal of Solids and Structures 62, 186–206 (2015).

50 Dong, H. et al. Micromechanical models for the stiffness and strength of UHMWPE macrofibrils. Journal of the Mechanics and Physics of Solids 116, 70–98 (2018).

51 Dong, H., Liu, M., Martin, C. & Sun, W. A residual stiffness-based model for the fatigue damage of biological soft tissues. Journal of the Mechanics and Physics of Solids 143, 104074 (2020).

52 Dong, H. & Hu, Y. Harnessing fluid pre-pressure to tune the properties of phononic crystals. Extreme Mechanics Letters 34, 100582 (2020).

53 Liu, M., Liang, L., Dong, H., Sun, W. & Gleason, R. L. Constructing growth evolution laws of arteries via reinforcement learning. Journal of the Mechanics and Physics of Solids, 105044 (2022).

54 Omojola, V. O. et al. Comparative Analysis of Arterial Compliance in Mice Genetically Null for Cathepsins K, L, or S. Journal of Biomechanics, 111266 (2022).

