## Supplementary Information for "Effect of Aging, Sex, and Gene (Fbln5) on Arterial Stiffness of Mice: 20 Weeks Adult Fbln5-knockout Mice Have Older Arteries than 100 Weeks Wild-Type Mice"

[3]Department *of Bioengineering*

Submitted to Some Journal

[a]These authors contributed equally.

[*]**For correspondence:**

Rudolph L. Gleason Jr, Ph.D.

The George W. Woodruff School of Mechanical Engineering
Georgia Institute of Technology
Technology Enterprise Park, Room 216
387 Technology Circle, Atlanta, GA 30313-2412

**S1. Theoretical pressure and non-reduced axial force based on the UFD model**

Under cylindrical biaxial testing condition, the through-thickness mean deformation gradient of the arterial tissue can be expressed as

$$\mathbf{F} = \mathrm{diag}[\lambda_r, \lambda_\theta, \lambda_z] \tag{SE1}$$

where $\lambda_r, \lambda_\theta,$ and $\lambda_z$ are stretches in the radius, circumferential and axial directions, respectively, with $\lambda_r = 1/(\lambda_\theta \lambda_z)$ by incompressibility. Then we have the invariants $\bar{I}_{4\theta} = \lambda_\theta^2$ and $\bar{I}_{4z} = \lambda_z^2$.

The Cauchy stress of the arterial tissue can be expressed as[1]

$$\boldsymbol{\sigma} = 2\mathbf{F}\frac{\partial\bar{\Psi}}{\partial\bar{\mathbf{C}}}\mathbf{F}^T - p\mathbf{I} \tag{SE2}$$

where $p$ is a Lagrange contribution to the hydrostatic pressure, and

$$\frac{\partial\bar{\Psi}}{\partial\bar{\mathbf{C}}} = \bar{\Psi}_1\mathbf{I} + \bar{\Psi}_4\mathbf{a}_\theta \otimes \mathbf{a}_\theta + \bar{\Psi}_{4\perp}\mathbf{a}_z \otimes \mathbf{a}_z \tag{SE3}$$

where

$$\bar{\Psi}_1 = \frac{\partial\bar{\Psi}}{\partial\bar{I}_1}, \bar{\Psi}_{4\theta} = \frac{\partial\bar{\Psi}}{\partial\bar{I}_{4\theta}}, \bar{\Psi}_{4z} = \frac{\partial\bar{\Psi}}{\partial\bar{I}_{4z}} \tag{SE4}$$

Substituting Eq. (2) of the main text into Eq. (SE4), we have

$$\bar{\Psi}_1 = \frac{c}{2},$$

$$\bar{\Psi}_{4\theta} = \delta_\theta k_1 \zeta^2(\bar{I}_{4\theta} - 1) \cdot \exp\{k_2[\delta_\theta\zeta^2(\bar{I}_{4\theta} - 1)^2 + \delta_z(1-\zeta)^2(\bar{I}_{4z} - 1)^2]\},$$

$$\bar{\Psi}_{4z} = \delta_z k_1 (1-\zeta)^2(\bar{I}_{4z} - 1) \cdot \exp\{k_2[\delta_\theta\zeta^2(\bar{I}_{4\theta} - 1)^2 + \delta_z(1-\zeta)^2(\bar{I}_{4z} - 1)^2]\}. \tag{SE5}$$

Substituting $\bar{I}_{4\theta} = \lambda_\theta^2$ and $\bar{I}_{4z} = \lambda_z^2$ into Eq. (SE5) and then into Eqs. (SE2)- (SE4), together with the condition of $\sigma_{rr} = 0$, the non-zero components of the Cauchy stress can be expressed as

$$\sigma_{\theta\theta} = c\left(\lambda_\theta^2 - \frac{1}{\lambda_\theta^2\lambda_z^2}\right) + 2\delta_\theta k_1 \lambda_\theta^2 \zeta^2(\lambda_\theta^2 - 1)\exp\{k_2[\delta_\theta\zeta^2(\bar{I}_{4\theta} - 1)^2 + \delta_z(1-\zeta)^2(\bar{I}_{4z} - 1)^2]\}$$

$$\sigma_{zz} = c\left(\lambda_z^2 - \frac{1}{\lambda_\theta^2\lambda_z^2}\right) + 2\delta_z k_1 \lambda_z^2 (1-\zeta)^2(\lambda_z^2 - 1)\exp\{k_2[\delta_\theta\zeta^2(\bar{I}_{4\theta} - 1)^2 + \delta_z(1-\zeta)^2(\bar{I}_{4z} - 1)^2]\} \quad \text{(SE6)}$$

Then the theoretical pressure $P_{model}$ and non-reduced axial force $f_{model}$ can be calculated based on the following equations

$$P_{model} = \frac{h\,\sigma_{\theta\theta}}{r_i}, \text{and } f_{model} = \pi(r_o^2 - r_i^2)\sigma_{zz} \quad \text{(SE7)}$$

where $h$ is the tissue thickness in loaded configuration. $r_i$ and $r_o$ are inner and out radius of the arterial tube in loaded configuration.

**S2. Stress under equal biaxial tension in axial (z) and circumferential ($\boldsymbol{\theta}$) directions**

Without loss of generality, we used the stress in the circumferential direction ($\sigma_{\theta\theta}$) to illustrate the physical meanings of $k_1$ $k_2$, $c$, and $\zeta$. Under equal biaxial tension with $\lambda_\theta = \lambda_z = \lambda > 1$, we have $\delta_\theta = \delta_z = 1$. The Cauchy stress $\sigma_{\theta\theta}$ can be further expressed as

$$\sigma_{\theta\theta} = c\left(\lambda^2 - \frac{1}{\lambda^4}\right) + 2k_1\lambda^2\zeta^2(\lambda^2 - 1)\exp\left\{k_2[(1-\zeta)^2 + \zeta^2](\lambda^2 - 1)^2\right\},$$

$$\sigma_{zz} = c\left(\lambda^2 - \frac{1}{\lambda^4}\right) + 2k_1\lambda^2(1-\zeta)^2(\lambda^2 - 1)\exp\{k_2[(1-\zeta)^2 + \zeta^2](\lambda^2 - 1)^2\}. \quad \text{(SE8)}$$

where the terms with $c$ is the contribution of the matrix, noted by

$$\sigma_{\theta\theta m} = \sigma_{zzm} = c\left(\lambda^2 - \frac{1}{\lambda^4}\right) \quad \text{(SE9)}$$

and the terms with $k_1$, $k_2$ and $\zeta$ is the contribution of the fiber, noted by

$$\sigma_{\theta\theta f} = 2k_1\lambda^2\zeta^2(\lambda^2 - 1)\exp\{k_2[(1-\zeta)^2 + \zeta^2](\lambda^2 - 1)^2\}$$

$$\sigma_{zzf} = 2k_1\lambda^2(1-\zeta)^2(\lambda^2 - 1)\exp\{k_2[(1-\zeta)^2 + \zeta^2](\lambda^2 - 1)^2\} \qquad \text{(SE10)}$$

The physical meaning of $k_1$ $k_2$, $c$, and $\zeta$ can be found in Fig. 6 of the main text. When fixing $\zeta$=0.5, for the fiber stress-stretch ($\sigma_{\theta\theta f} - \lambda_\theta$) curve with different $k_2$ but fixed $k_1$ (Fig. 6a), the initial slope (initial modulus) does not change, while the stiffening effect increases with increasing $k_2$. In Fig. 6b (different $k_1$ but fixed $k_2$), the initial slope (i.e., initial modulus) of the fiber stress-stretch ($\sigma_{\theta\theta f} - \lambda_\theta$) curve increases proportionally with increasing $k_1$, while the stiffening effect, characterized by the ratio between the stiffening tangent modulus and the initial tangent modulus, does not change. The parameter $c$ controls the stiffness of the matrix, which has little stiffening effect at the large deformation (Fig. 6c). Thus, the parameter $k_2$ determines stiffening of the entire tissue (matrix has little stiffening effect). Fig. 6d shows how $\zeta$ (ranging from 0 to 1) influences the ratio $2\sigma_{\theta\theta f}/(\sigma_{\theta\theta f} + \sigma_{zzf})$, which indicates that $\zeta$ controls the degree of anisotropy between axial and circumferential directions.

### References

1 Holzapfel, G. A. *Nonlinear solid mechanics: A continuum approach for engineering science*. (John Wiley & Sons, Ltd, 2000).
